# Pharmacologic inhibition of BAF chromatin remodeling complexes as a therapeutic approach to transcription factor-dependent cancers

**DOI:** 10.1101/2023.09.11.557162

**Authors:** Richard C. Centore, Luis M. M. Soares, Salih Topal, Rishi G. Vaswani, Kana Ichikawa, Zhifang Li, Hong Fan, Jeremy W. Setser, David L. Lahr, Laura E. Zawadzke, Xueying Chen, Kimberly D. Barnash, Jordana Muwanguzi, Neville Anthony, Gabriel J. Sandoval, Katharine Feldman, GiNell Elliott, Ammar Adam, David Huang, Yunji Davenport, Shawn Schiller, Kevin J. Wilson, Johannes Voigt, Lan Xu, Martin Hentemann, David S. Millan, Ho Man Chan, Carl P. Decicco, Ryan G. Kruger, Gromoslaw A. Smolen, Steven F. Bellon

**Affiliations:** Foghorn Therapeutics, 500 Technology Square, Suite 700, Cambridge, MA 02139

## Abstract

The BRG/Brahma-associated factors (BAF or mSWI/SNF) family of chromatin remodeling complexes are critical regulators of gene expression and are major determinants of cancer and other diseases. Two paralog ATPases, SMARCA4 and SMARCA2 (BRG1 and BRM, respectively), provide the enzymatic activity required for chromatin remodeling. Here, we discover and characterize a novel series of compounds that potently and selectively inhibit SMARCA4/SMARCA2. Mutational and biochemical studies demonstrate that these inhibitors act through a unique mode of inhibition, distinct from reported SMARCA4/SMARCA2 inhibitors. Across a range of cancer cell lines, SMARCA4/SMARCA2 inhibition resulted in lineage-specific changes in chromatin accessibility at binding sites for key transcription factors (TFs). In uveal melanoma (UM), BAF inhibition resulted in loss of enhancer occupancy of SOX10 and MITF, two essential TFs, leading to down-regulation of the melanocytic gene expression program. In a mouse xenograft model of UM, SMARCA4/SMARCA2 inhibition was well tolerated and resulted in dose-dependent tumor regression correlating with pharmacodynamic modulation of BAF-target gene expression. These data provide the foundation for first-in-human studies of BAF ATPase inhibition as a novel therapeutic to treat TF-dependent cancers.

---

Targeting transcriptional regulators has emerged as an exciting therapeutic approach in oncology. In recent years, inhibitors of histone or DNA methyltransferases (*e*.*g*., EZH2), histone demethylases (*e*.*g*., LSD1), histone deacetylases, histone modification readers (*e*.*g*., BRD4), and transcription-associated kinases (*e*.*g*., CDK7), have advanced to clinical development or, in the case of the EZH2 inhibitor, Tazemetostat, have received FDA approval^1,2^. Chromatin remodeling is another critical aspect of transcriptional regulation that holds promise as an additional means for disease intervention. Several large, multi-protein complexes including BAF, ISWI, NuRD and INO80 use the energy generated by ATP hydrolysis to remodel chromatin^3^. Of these, the BAF family of chromatin remodeling complexes has the strongest link to human cancer, as many of its subunits are mutated in a disease-specific manner, providing unique tumor vulnerabilities^4^. To date, three distinct BAF complexes have been identified: canonical BAF (cBAF), polybromo-associated BAF (PBAF), and non-canonical BAF (ncBAF)^5,6^. Each of these BAF complexes incorporates, at its catalytic core, one of two mutually exclusive ATPases, SMARCA4 or SMARCA2^7–9^. Through ATP-dependent chromatin remodeling, BAF complexes are responsible for establishing and maintaining tissue- and disease-specific chromatin landscapes conducive to specific transcriptional outcomes^10–14^. Dysregulation of BAF activity is implicated in various disease mechanisms, with mutations in genes encoding components of the BAF complexes most prevalent in cancer and neurological disorders^15^.

Targeting the ATPase activity of BAF may provide a promising therapeutic approach in oncology. Functional genomics screens using shRNA or CRISPR have revealed dependencies on SMARCA4 or SMARCA2 in many cancer cell lines with diverse genomic contexts^16–18^. These two proteins have a synthetic lethal relationship. Genetic depletion of *SMARCA2* by shRNA or CRISPR results in selective growth inhibition of cancer cell lines harboring loss-of-function mutations in *SMARCA4*^19–21^. Recently, this relationship was shown to hold true in the opposite direction as well, as genetic and pharmacological targeting of *SMARCA4* was shown to result in selective growth inhibition of esophageal cancer cell lines with low *SMARCA2* expression as compared to those with high *SMARCA2* levels^22^. Given the duality of this relationship, coupled with the extensive percentage of human cancer cell lines that are dependent on BAF complexes, we determined that a small molecule acting as a dual inhibitor of SMARCA4 and SMARCA2 ATPases could have utility in the treatment of a broad spectrum of cancer indications.

To explore the therapeutic potential of SMARCA4/SMARCA2 ATPase inhibition, we carried out a high-throughput screening campaign which led to the discovery of a novel chemical series that potently inhibits both SMARCA4 and SMARCA2, while sparing other related ATPases with highly similar ATPase domains. This series is structurally distinct from a recently disclosed SMARCA4/SMARCA2 tool compound, and inhibits SMARCA4/SMARCA2 through a different mode of action^23^. With our dual SMARCA4/SMARCA2 inhibitor we show that BAF inhibition alters the chromatin landscape by attenuating enhancer accessibility at transcription factor binding sites in cancer cell lines from multiple lineages. Using uveal melanoma as a model, we further demonstrate that *in vitro* and *in vivo* abrogation of lineage transcription factor function through pharmacologic inhibition of BAF ATPases offers a novel therapeutic approach to treat cancers addicted to these transcriptional networks. Thus, while selective SMARCA2 inhibition is an attractive strategy for exploiting synthetic lethal relationships, our findings suggest that dual SMARCA4/SMARCA2 ATPase inhibition could provide a therapeutic means to target cancers that are exquisitely dependent upon BAF-cooperating transcription factors.

## Results

### Discovery of a novel class of SMARCA4/SMARCA2 ATPase inhibitors

To identify ATPase inhibitors of SMARCA4/SMARCA2, we screened a 165,000 compound library for inhibitors of DNA-dependent ATPase activity of full length SMARCA2. The initial screening hit, FHT-185, demonstrated half-maximal inhibitory concentration (IC_50_) of 2.5 ± 0.5 µM (n=7), and represented a novel chemical scaffold for this target class (Fig. 1a-b). Optimization of this scaffold for potency through traditional medicinal chemistry efforts led to the identification of the *in vitro* tool compound, FHT-1015; and additional optimization for ADME (absorption, distribution, metabolism and excretion) properties led to the identification of FHT-2344. Both FHT-1015 and FHT-2344 were characterized as dual SMARCA4/SMARCA2 ATPase inhibitors, with biochemical IC_50_ values in the low nanomolar range for the two targets (Fig. 1b-c). Notably, neither molecule showed inhibition of the related SNF2-family ATPase,CHD4, up to concentrations > 200 µM (Fig. 1b, Supplementary Fig. 1a). In a cell-based assay platform to measure competitive binding of compounds to ATP-binding pockets of hundreds of ATPases (KiNativ), FHT-2344 exhibited little binding to ATPases in general, demonstrating further selectivity (Supplementary Fig. 1b). Together, these data suggest that these compounds are potent and selective inhibitors of the SMARCA4/SMARCA2 ATPases.

**Figure 1:**
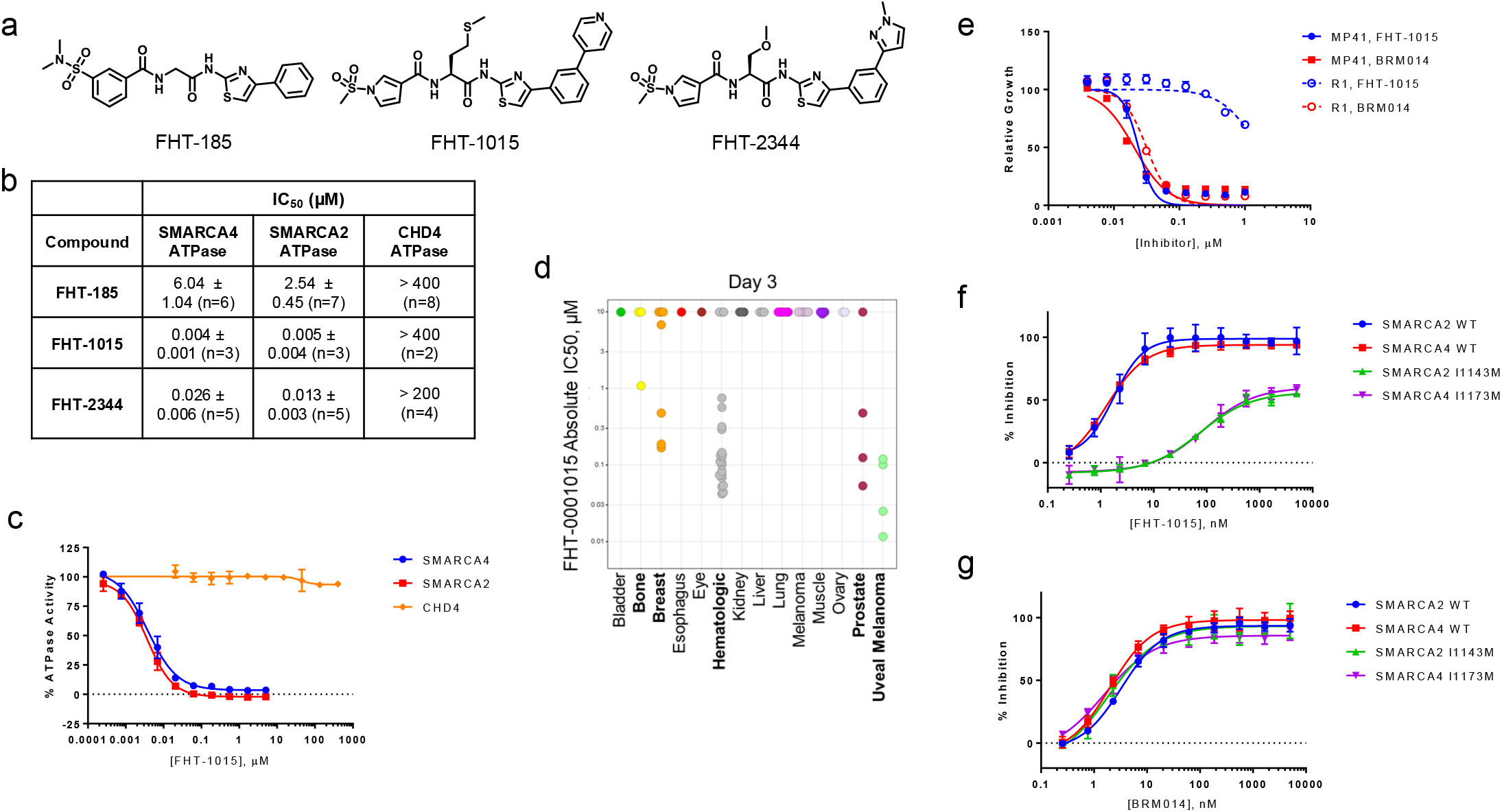
Identification and optimization of a novel class of dual inhibitors of SMARCA4/SMARCA2 ATPase activity. (**a**) Chemical structures of FHT-185, FHT-1015, and FHT-2344. (**b**) Selective inhibition ofSMARCA4 and SMARCA2 ATPases. DNA-dependent ATPase activity was measured by the ADP-glo assay in the presence of inhibitor using full length SMARCA4, SMARCA2 and CHD4. IC_50_ values for the 3 compounds from (a) (geometric mean ± SD) are shown. (**c**) Representative dose response curves from (b) (mean ± SD). (**d**) Effects of FHT-1015 on the proliferation of cell lines were tested in a 3-day assay by Cell-titer glo (CTG). Absolute IC_50_ values are plotted, and each dot represents an individual cell line. (**e**) Effects of FHT-1015 and BRM014 on the proliferation of parental MP41 or MP41-derived cells that are resistant to FHT-1015 (R1). Relative growth is plotted following a 3-day CTG assay (n=2; mean ± SD). (**f-g**) Inhibitory effects of FHT-1015 (**f**) or BRM014 (**g**) on wild type and mutant SMARCA4 or SMARCA2 ATPases in a Pyruvate Kinase/Lactate Dehydrogenase (PK/LDH) coupled enzyme assay (n = 3, mean ± SD).

### BAF ATPase inhibition affects the proliferation of cancer cell lines

*SMARCA4* is highly expressed across cancer subtypes, and functional genomics studies have demonstrated its dependency in many cancer cell lines, particularly those of hematologic, esophageal, prostate and uveal melanoma origin (Supplementary Fig. 1c)^16,17^. To test the impact of SMARCA4/SMARCA2 ATPase inhibition (also referred to hereafter as BAF ATPase inhibition or BAFi), we screened a small panel of cancer cell lines, with representation from each of these lineages, for sensitivity to FHT-1015 in a 3- or 7-day growth inhibition assay. While most cell lines screened had a robust growth inhibitory response to FHT-1015 at day 7, only a subset of cell lines demonstrated sub-micromolar response at the earlier, day 3 time-point (Fig. 1d, Supplementary Fig. 1d). These fast-responding cell lines may represent indications that are particularly sensitive to SMARCA4/SMARCA2 inhibition. Notably, cell lines from hematologic cancers and uveal melanoma were overrepresented in this group. We decided to focus on uveal melanoma (UM) given that every one of the cell lines of this lineage tested belonged to the most sensitive, fast-responding group.

### FHT-1015 inhibits SMARCA4/SMARCA2 through a novel mode of inhibition

To confirm on-target activity of FHT-1015, we first attempted to establish a UM model of acquired resistance to this compound. MP41 cells were grown in the presence of 0.1 µM FHT-1015 for about three months. At this concentration, greater than 95% of MP41 cell growth is inhibited within three days. After three months, FHT-1015-resistant clones were observed. These clones grew at the same rate as the parental MP41 cells, but were refractory to growth inhibition by FHT-1015 (Fig. 1e, Supplementary Fig. 2a). Importantly, the resistant cells were equally as sensitive as the parental MP41 cells to treatment by IDE196 (PKC inhibitor) or paclitaxel, demonstrating that the resistance mechanism is highly specific and not due to a general increase in fitness (Supplementary Fig. 2b). To identify resistance-conferring mutations, we next subjected 9 resistant isolates as well as the parental MP41 cells to whole exome sequencing (WES). Interestingly, all 9 resistant isolates harbored a common heterozygous SMARCA4 mutation: I1173M (Supplementary Fig. 2c). To test if this mutation was indeed resistance-conferring, we generated recombinant SMARCA4 with the I1173M mutation, and SMARCA2 with the homologous I1143M mutation, and tested them in parallel with wild type SMARCA4 or SMARCA2 enzymes in an enzymatic ATPase assay. Indeed FHT-1015 demonstrated significantly reduced inhibition of the SMARCA4 I1173M and SMARCA2 I1143M mutants (Fig. 1f). Therefore, this single amino acid mutation renders SMARCA4/SMARCA2 resistant to FHT-1015, supporting the notion that SMARCA4 is the primary target in FHT-1015-mediated growth inhibition of MP41.

We noted that BRM014, a previously reported SMARCA4/SMARCA2 inhibitor^23^, was equally effective in inhibiting the parental and FHT-1015-resistant MP41 cell proliferation (Fig. 1e). BRM014 also inhibited the ATPase activities of wild type SMARCA4/SMARCA2 and mutant I1173M/I1143M enzymes with equal potency (Fig. 1g). Interestingly, I1143 of SMARCA2 (homologous to I1173 of SMARCA4) resides outside of the N-terminal RecA lobe of SMARCA2 (amino acids 705-960) which was shown to be the binding sitefor BRM014^23^. Together, these data demonstrate that FHT-1015 is a novel on-target inhibitor of SMARCA4 and SMARCA2 ATPases, with a mode of inhibition distinct from that of BRM014.

### SMARCA4/SMARCA2 inhibition induces apoptosis in uveal melanoma cells

Given that 100% of the UM cell lines tested exhibited significant growth inhibition upon treatment with FHT-1015, we pursued further mechanistic characterization in this disease context. First, to further understand how SMARCA4/SMARCA2 inhibition suppresses UM cell proliferation, we examined a panel of 4 uveal melanoma derived cell lines and compared their growth inhibition profiles to that of the *SMARCA4* mutant non-small cell lung cancer (NSCLC) cell line, NCIH1299. All 4 UM cell lines (92-1, MP38, MP41 and MP46) had an absolute IC_50_ of less than 0.1 µM, while NCIH1299 cells did not reach this threshold at any dose after 3 days of treatment (Supplementary Fig. 3a). This rapid activity of FHT-1015 suggested that UM cell lines were undergoing a different type of response from NSCLC cells upon BAF inhibition. Cell cycle profiling of 92-1 UM cells treated with FHT-1015 revealed an early G1 arrest followed by accumulation of cells in the sub-G1 state, suggesting UM cell apoptosis (Supplementary Fig. 3b-c). This phenotype differs from the previously observed G1 arrest and senescence of *SMARCA4-* mutant NSCLC cell lines upon genetic depletion of *SMARCA2*^19,20^. In UM cell lines, treatment with higher doses of FHT-1015 induced strong Caspase 3/7 activity and Annexin V staining, further supporting an apoptotic response of this cell type to BAF ATPase inhibition (Supplementary Fig. 3d-f). Together, these data suggest that uveal melanoma cell lines respond to SMARCA4/SMARCA2 inhibition through an apoptotic mechanism.

Uveal melanoma is a rare cancer of the eye with relatively low mutational burden, but in over 90% of cases, is driven by gain-of-function activating mutations in G_α_ signaling proteins (encoded by *GNAQ* or *GNA11*), which transduce proliferative signals, in part, through Protein Kinase C (PKC) and MAPkinases^24,25^. Inhibitors of PKC and MEK are currently in clinical development for the treatment of uveal melanoma, although to date, with very limited clinical success^26–28^. Comparison of FHT-1015 to IDE196 or Selumetinib, inhibitors of PKC or MEK, respectively, revealed that FHT-1015 was more potent than each of these compounds in a 3-day proliferation assay, with absolute IC_50_ shifts of at least one order of magnitude in both 92-1 and MP41 UM cell lines (Supplementary Fig. 3g-h). Consistent with its slower growth inhibitory effect, IDE196 did not induce apoptosis as measured by cell cycle distribution, Caspase 3/7 activity, or Annexin V staining (Supplementary Fig. 3b-c,e-f,i). Thus, BAF inhibition represents a novel modality with a different phenotypic effect from other compounds being tested in the clinic for the treatment of uveal melanoma.

### SMARCA4/SMARCA2 inhibition alters enhancer accessibility of SOX10 and other disease-relevant transcription factors

To determine the global effects of BAFi on chromatin accessibility in UM, we performed the Assay for Transposase-Accessible Chromatin coupled with next generation sequencing (ATAC-seq) in the 92-1 cell line. FHT-1015 treatment did not result in genome-wide loss of accessibility at all sites detectable by ATAC-seq, but rather had a specific effect at a subset of sites (Fig. 2a, Supplementary Fig. 4a). ATAC-seq peaks were binned into three clusters: Cluster 1, loss of accessibility peaks; Cluster 2, peaks with no change in accessibility; and Cluster 3, gain of accessibility peaks, in response to FHT-1015 treatment. BAF inhibition generally resulted in closing of chromatin, as 13,698 Cluster 1 peaks existed with reduced accessibility and only 58 Cluster 3 peaks were identified with increased accessibility following FHT-1015 treatment (Fig. 2a, Supplementary Fig. 4a-b). The Cluster 1 regions of reduced accessibility upon FHT-1015 treatment occurred at sites of dense SMARCA4-enrichment as measured by Chromatin Immuno-Precipitation followed by next-generation sequencing (ChIP-seq) (Fig 2a, Supplementary Fig. 4b). Given that BAF complexes can regulate chromatin accessibility at both gene promoters and distal regulatory enhancer elements^29^, we further classified ATAC-seq peaks as promoter- or enhancer-occupied using annotatePeaks function of HOMER package^30^. Strikingly, FHT-1015 treatment resulted in a marked loss of accessibility at enhancers, with only a modest effect at promoters (Fig. 2b, Supplementary Fig. 4c). Motif analysis at Cluster 1 regions (loss of accessibility peaks) revealed the enrichment of several transcription factor binding sites, including SOX10, TFAP2A, and to a lesser degree MITF (Fig. 2c). SOX10, TFAP2A, and MITF are essential for the development and maintenance of melanocytes, a cell type derived from neural crest^31-35^. Furthermore, all three transcription factors were among the top-ranked gene dependencies in UM cell lines with *GNAQ* or *GNA11* mutations; and their ranking in gene dependency was comparable to that of the disease driver G_α_ as well as *SMARCA4* itself (Supplementary Fig. 4d).

**Figure 2:**
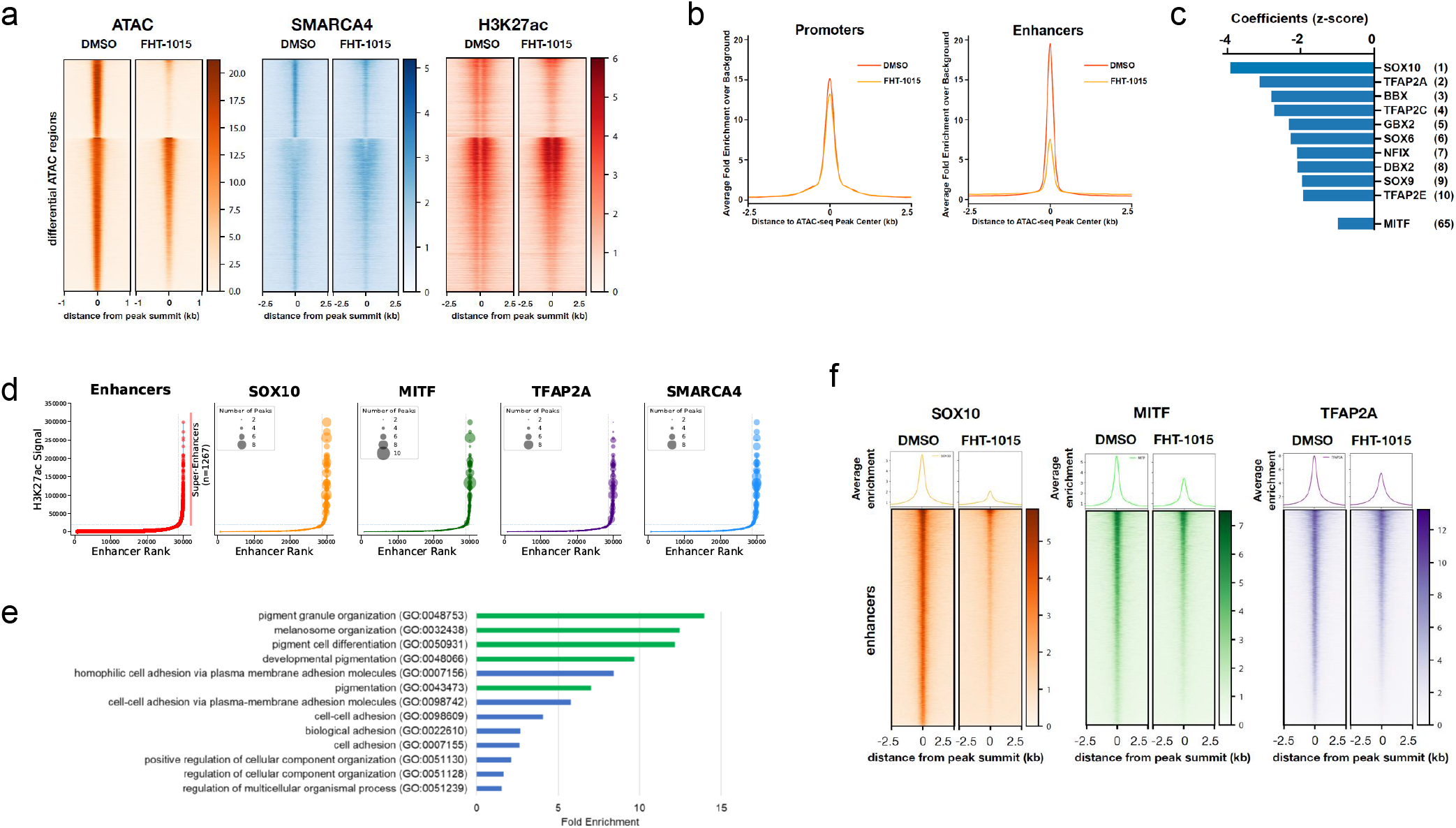
BAF ATPase inhibition alters enhancer accessibility of disease-relevant transcription factors. (**a**) 92-1 cells were treated for 4 hours with DMSO or 100 nM FHT-1015. Heatmaps with normalized coverage tracks for DMSO or FHT-1015 treated samples are shown for all chromatin accessible regions identified in 92-1 cells by ATAC-seq. Regions were classified as consistently losing accessibility, not changing, or gaining accessibility, and sorted according to signal in the DMSO treated sample. Average signal for ATAC-seq, SMARCA4 ChIP-seq and H3K27Ac ChIP-seq was calculated and depicted as heatmaps centered at the ATAC-seq peak maximum value. Flanking region from the center of the ATAC-seq peak is 1 kb for ATAC-seq heatmap and 2.5 kb for SMARCA4/H3K27Ac ChIP-seq heatmaps. (**b**) Chromatin accessible regions depicted in (a) were annotated as promoter- or enhancer-associated. For each, group average signal for ATAC-seq was calculated and depicted as anchor plots centered at the ATAC-seq peak maximum value (**c**) Transcription factor motifs enriched at sites of lost accessibility from (a). For each region of accessible chromatin identified, potential transcription factor binding sites were assigned. The correlation between the presence of a transcription factor binding site and the observed change in chromatin accessibility upon FHT-1015 treatment was modelled using ridge regression, thecoefficients of the regression were z-scored and plotted. The presence of SOX10 motifs shows the most negative coefficient indicating the stronger effect in explaining the loss of accessibility observed. (**d**) 92-1 super-enhancers are enriched for SOX10, MITF, TFAP2A, and SMARCA4 binding. For each enhancer/super-enhancer identified in 92-1 cells, the number of ChIP-seq peaks of the indicated factor was calculated and plotted. (**e**) Gene ontology (GO) of the top 500 genes mapped to 92-1 super-enhancers identified in (d) was run using PANTHER overrepresentation test. Fold enrichment for GO biological process categories with a false discovery rate (FDR) < 0.05 in a FISHER test are plotted. (f) Average ChIP-seq enrichment of SOX10, MITF, and TFAP2A at enhancers following treatment of 92-1 cells with DMSO or 100 nM FHT-1015 for 4 hours. Anchor plots are centered on ChIP-seq peaks for each indicated TF.

### SMARCA4/SMARCA2 inhibition modifies lineage-defining enhancer landscapes

Consistent with an important role for SOX10, TFAP2A, and MITF in UM biology, ChIP-seq in 92-1 cells showed enrichment of all three TFs along with SMARCA4 at super-enhancers, as defined by promoter-distal regions with disproportionately high H3K27ac ChIP-seq signal (Fig. 2d). These UM super-enhancers were associated with genes involved in pigmentation and melanosome organization, key processes of melanocyte biology (Fig. 2e). Genetic depletion of *SMARCA4* has been shown to cause preferential loss of H3K27ac at enhancers as compared to promoters^14^. Indeed, pharmacologic inhibition of SMARCA4/SMARCA2 with FHT-1015 resulted in a similar effect, with reduced H3K27ac levels observed at a subset of enhancers, including super-enhancers at important melanoma genes, *SOX10, MITF*, and *PMEL* (Supplementary Fig. 4e). In contrast, at Cluster 2 sites with no change in accessibility, including at promoters, H3K27ac was markedly increased after FHT-1015 treatment, andSMARCA4 occupancy sites became more broad (Supplementary Fig. 4b). The reason for this accumulation is not clear, although it may reflect a feedback mechanism or a disconnect in enhancer-promoter communication. Together, these data point to the ATPase activity of BAF as a critical regulator of lineage-defining enhancer structures.

### BAF ATPase activity regulates transcription factor binding at enhancers

The enrichment of SOX10, TFAP2A, and MITF motifs at sites of BAF-regulated accessibility, as well as the co-occupancy of SMARCA4 with these TFs at super-enhancers suggested that BAF ATPase activity may be required to allow access of these TFs to their binding sites. To directly test this idea, we analyzed the effects of BAFi on SOX10, MITF, and TFAP2A enhancer occupancy by ChIP-seq. Upon a 4-hour treatment with FHT-1015, enhancer occupancy of each of these TFs was markedly reduced, with no effect on their overall protein abundance at this timepoint, suggesting that the ATPase activity of BAF complexes is required for these transcription factors to access their genomic binding sites (Fig. 2f, Supplementary Fig. 4f). The loss of chromatin occupancy of each of these TFs correlated well with the loss of accessibility induced by FHT-1015 treatment (Supplementary Fig. 4g-i). Further, Precision Nuclear Run-On (PRO)-seq in the MP46 UM cell line showed that SOX-family and MITF transcription factor binding motifs were enriched at transcription start sites of nascent transcripts that were down-regulated quickly (15 min – 1 hour) upon BAF inhibition (Supplementary Fig. 4j). Together, these data suggest that BAF ATPase inhibition results in rapid loss of chromatin accessibility at the binding sites for important, disease-relevant transcription factors, and prevents the binding of these TFs to chromatin.

### SMARCA4/SMARCA2 inhibition disrupts the SOX10-MITF transcriptional axis in uveal melanoma

To understand the impact of enhancer dysregulation caused by SMARCA4/SMARCA2 ATPase inhibition, we next examined the transcriptional consequences of FHT-1015. RNA-seq of 92-1 cells treated with FHT-1015 revealed 833 genes down-regulated, while only 386 were up-regulated 2-fold or more after a 4-hour treatment (Fig. 3a, Supplementary Fig. 5a). The expression profile upon BAF ATPase inhibition was very different from that of PKC inhibition, supporting the notion that these two modalities act through distinct mechanisms (Supplementary Fig. 5a). Gene set enrichment analysis (GSEA) identified enrichment of gene sets associated with SMARCA4 targets, melanoma, pigmentation, and neural precursor among genes down-regulated by BAF ATPase inhibition, consistent with the reduced binding of SOX10 and MITF to enhancers (Fig. 3b-c, Supplementary Fig. 5b). This relationship was reinforced when we examined the convergence of genes that were associated with SMARCA4 and H3K27ac occupancy, reduced accessibility, and loss of expression upon FHT-1015 treatment, as 53 genes met these criteria and were enriched for gene ontology terms associated with melanoma and pigmentation processes (Fig. 3d). Down-regulation of SOX10/MITF target genes was extended to additional UM cell lines (MEL202, MP41 and 92-1) as well (Fig. 3e). Biological pathways that were up-regulated by FHT-1015 included multiple neuron differentiation and developmental pathways, including: negative regulation of neural crest differentiation, consistent with known biological functions of SOX10 (Supplementary Fig. 5c). Like many other lineage-specifying transcription factors, SOX10 has been shown to regulate its own expression^36^. Indeed, in 92-1 cells, we observed occupancy of SOX10, MITF, TFAP2A and SMARCA4 at the *SOX10* super-enhancer, marked by accessible chromatin and H3K27ac (Fig. 3f). Remarkably, each of these factors showed reduced binding at the *SOX10* enhancers following a 4-hour treatment with FHT-1015 (Fig. 3f). Concurrently, *SOX10* expression was significantly down-regulated in 3 different UM cell lines (MEL202, MP41 and 92-1) (Fig. 3a,e-f, Supplementary Fig. 5d-e). Further, PRO-seq demonstrated that nascent transcripts at the *SOX10* gene body and enhancer were strongly reduced after just 1 hour of FHT-1015 treatment, suggesting that *SOX10* gene suppression is among the first transcriptional events to occur upon BAF ATPase inhibition (Fig. 3g-h, Supplementary Fig. 5f). In two UM cell lines (92-1 and MEL202), *SOX10* clearly stood out as a gene with a strong dependency score and with robust BAFi-induced down-regulation (Fig. 3i, Supplementary Fig. 5g). In comparison, the expression of *MITF* was also reduced, but the kinetics of *MITF* down-regulation were markedly slower than observed for *SOX10* (Fig. 3i, Supplementary Fig. 5d,g). These data suggested that *SOX10* may be the most upstream dependency node in UM.

**Figure 3:**
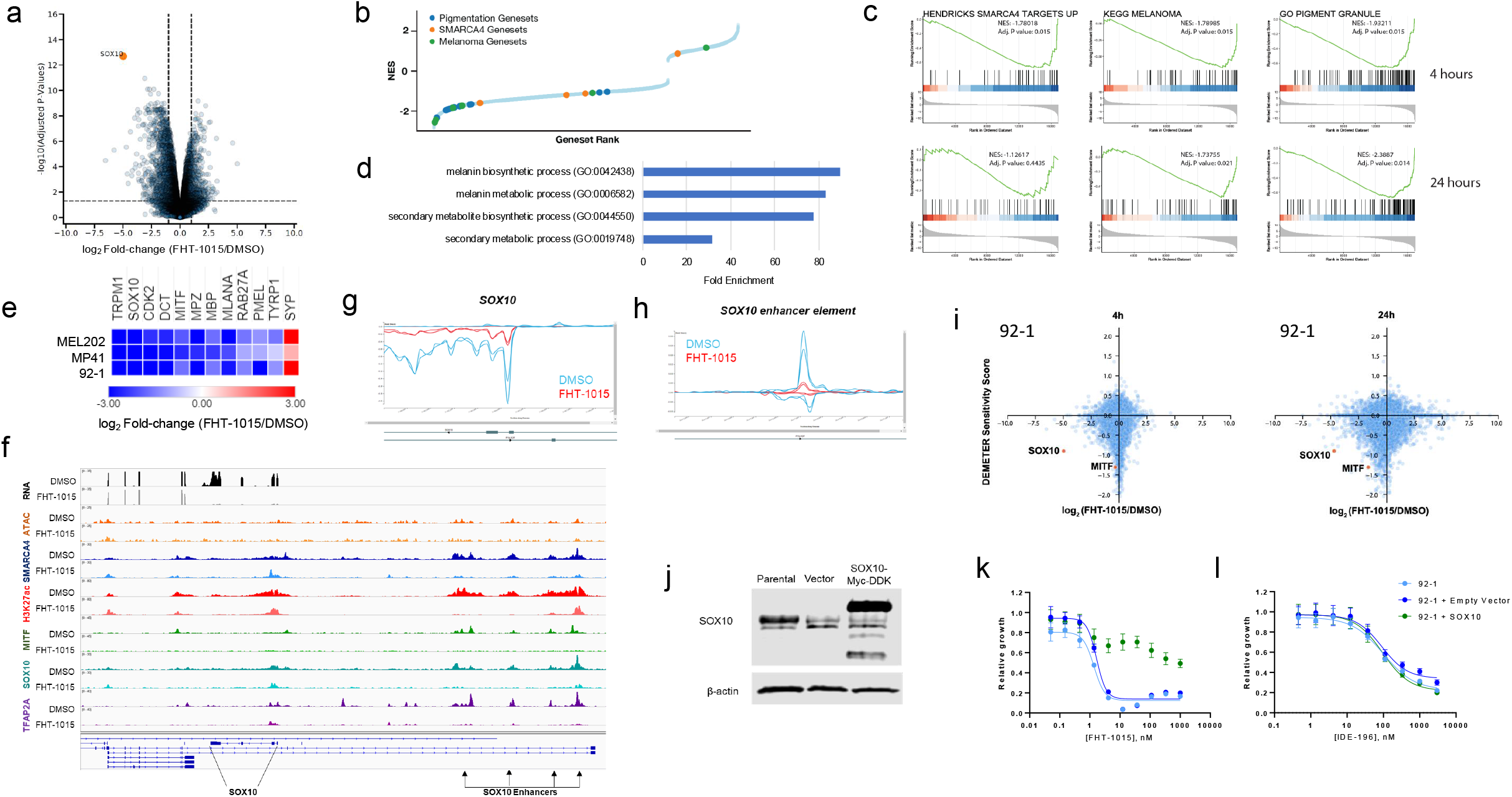
BAF ATPase inhibition disrupts the SOX10-MITF transcriptional axis in uveal melanoma. (**a**) Volcano plot depicting results of differential gene expression analysis of 92-1 cells treated with DMSO or FHT-1015 after 4 hours. SOX1*0* differential expression is highlighted in orange. Vertical dashed lines indicate 2-fold expression change thresholds; horizontal dashed line indicates adjusted p-value of 0.05. (**b**) Gene set enrichment analyses (GSEA) of 92-1 cells treated with DMSO or 100 nM FHT-1015 for 24h. Differential gene expression effects were ranked according to adjusted p-value and signal of change, and the ranked list was used for GSEA using MSIGDB gene sets. Enrichment of pigmentation, melanoma, and SMARCA4 target gene sets among those down-regulated by FHT-1015 is highlighted. (**c**) Representative GSEA plots of gene sets that are enriched among genes down-regulated by FHT-1015 in 92-1 cells. (**d**) Panther gene ontology enrichment of melanoma/pigmentation pathway for 53 genes characterized by loss of accessibility after a 4h treatment with FHT-1015 (ATAC-seq log_2_FC < −1), loss of expression after 24h treatment with FHT-1015 (RNA-seq log_2_FC < −3), and occupancy of SMARCA4 and H3K27ac by ChIP-seq in 92-1 cells (**e**) Heatmap representing SOX10/MITF target genes that are consistently modulated by FHT-1015 across 3 uveal melanoma cell lines. Cells were treated with DMSO or 100 nM FHT-1015 for4h. (**f**) Genome browser view of RNA-seq, ATAC-seq, and ChIP-seq of SMARCA4, H3K27ac, and transcription factors at the *SOX10* locus in 92-1 cells treated for 4 hours with DMSO or 100 nM FHT-1015. (**g-h**) PRO-seq tracks at the *SOX10* gene body (**g**) or enhancer (**h**) in MP46 cells following a 1 hour treatment of DMSO or 100 nM FHT-1015. (**i**) 92-1 cells were treated with DMSO or 100 nM FHT-1015 for 4h or 24h and gene expression changes were plotted against the DEMETER dependency score^46^. *SOX10* is highlighted as a strong dependency with rapid and robust suppression of gene expression upon FHT-1015 treatment. (**j**) 92-1 cells were transduced with lentivirus with constitutive expression of empty vector or *SOX10-Myc-DDK*, and Western blot showing SOX10 protein expression after 7 days is shown alongside that of the parental 92-1 cells. (**k**) Resistance to FHT-1015-induced growth inhibition in 92-1 cells over-expressing *SOX10* in a 3-day CTG assay. (**l**) 92-1 cells over-expressing *SOX10* remain sensitive to IDE-196 in a 3-day CTG assay.

### Forced expression of SOX10 can overcome the phenotypic effects of SMARCA4/SMARCA2 inhibition

To further test if SOX10 downregulation by BAFi is a critical event in suppressing UM cell proliferation, we forced *SOX10* expression from a constitutive promoter and tested the effects on FHT-1015-dependent growth inhibition. Interestingly, *SOX10* overexpression resulted in a partial rescue of the phenotypic growth inhibition effects elicited by FHT-1015 (Fig. 3j-k). This rescue was specific to the growth inhibition caused by BAF ATPase inhibition, as SOX10 overexpression did not impact the ability of the PKC inhibitor, IDE-196, to cause growth inhibition of 92-1 cells (Fig. 3l). Together, these results suggest that SOX10 down-regulation contributes to the molecular mechanism underlying BAF ATPase inhibition in UM cell lines.

### SMARCA4/SMARCA2 inhibition causes uveal melanoma tumor regression in vivo

FHT-1015 provides a useful tool for SMARCA4/SMARCA2 ATPase inhibition *in vitro*, however the molecule lacks desired in vivo properties. Thus, to test the effects of BAF inhibition *in vivo*, we turned to FHT-2344, a related compound from the same series with improved ADME properties (Fig. 1a-b, Supplementary Fig. 6a). In a 21-day study using mice xenografted with the 92-1 UM cell line, FHT-2344demonstrated dose-dependent tumor growth inhibition (Fig 4a-b). Dosing at 2.2 mpk (mg/kg) daily (QD) led to 25% TGI while dosing at 6.7 mpk led to 92% TGI. Tumor regression was observed at the highest dose of 20 mpk. FHT-2344 was also well tolerated with no evidence of significant animal body weight loss in all dose groups (Fig. 4c). The observed anti-tumor efficacy correlated well with the suppression of both *SOX10* and a SMARCA4/SOX10 target gene, *FGF9*, as the pharmacodynamics biomarkers in the tumor (Fig. 4d-e, Supplementary Fig. 6b). Interestingly, as the plasma concentration of FHT-2344 decreased, the expression levels of *SOX10* appeared to rebound quickly to baseline (Fig. 4d). In contrast, FGF9, stayed suppressed to a greater extent at later timepoints (Fig. 4e). This observation could be attributable to differences in rates of transcription of the two genes or could point to compensation at the SOX10 locus, given the importance of this gene in maintaining UM survival. Together, these data underpin the importance of BAF in regulating *SOX10* expression as a mechanism of supporting UM tumor cell proliferation and survival *in vivo*.

**Figure 4:**
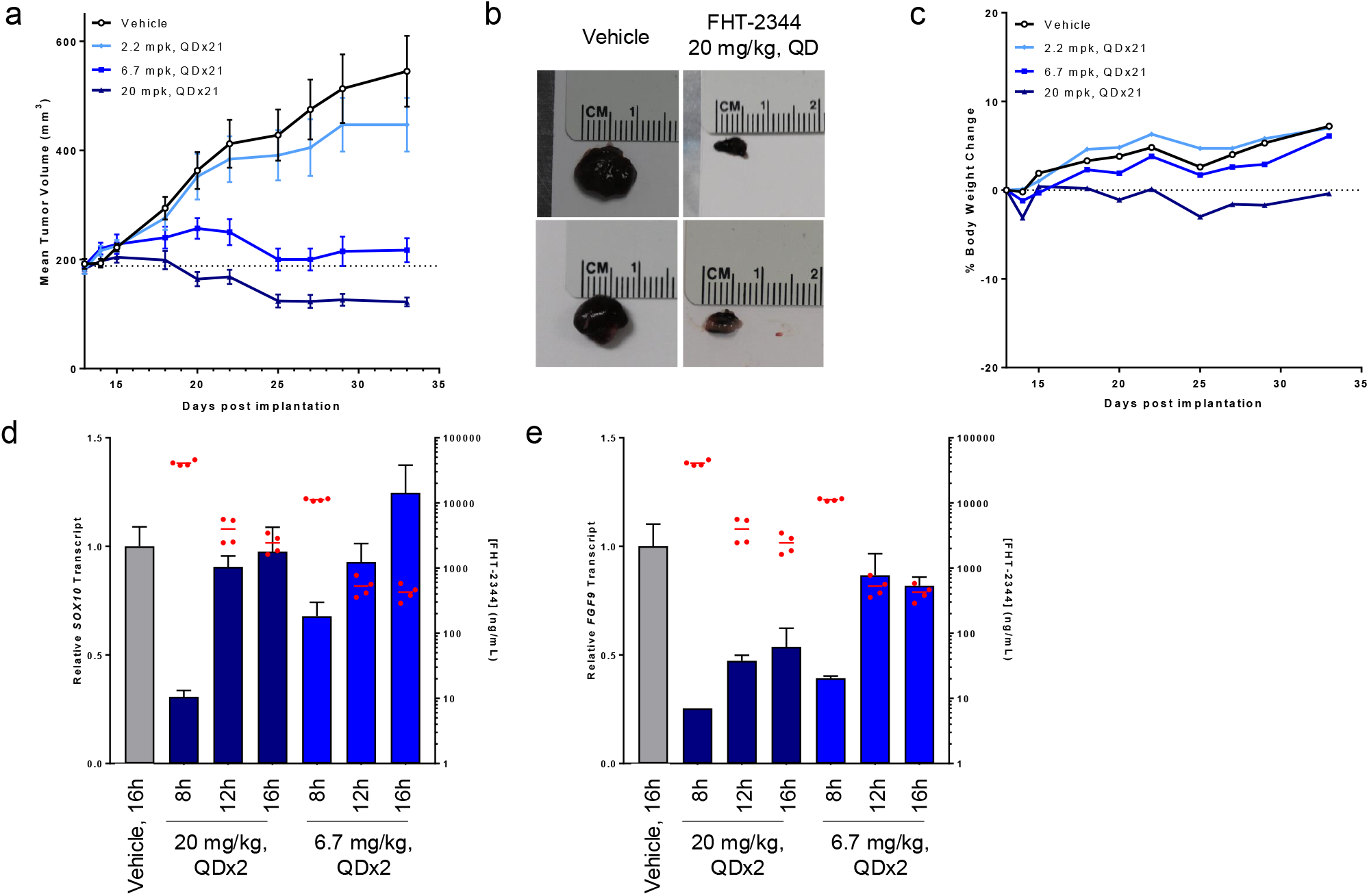
SMARCA4/SMARCA2 ATPase inhibition causes uveal melanoma tumor regression. (**a**) 92-1 cells were xenografted into nude mice and the effects of daily oral dosing of vehicle or FHT-2344 are plotted (n=10, mean ± SEM). (**b**) Representative images of tumors from (a) after 21 days of treatment. (**c**) Percent body weight change from (a) (n=10, mean ± SEM). (**d-e**) Pharmacokinetic and pharmacodynamic response of FHT-2344 in nude mice harboring 92-1 tumor xenograft. Following two days of once-daily, oral dosing, FHT-2344 was measured in the plasma (red dots, right axis), and relative gene expression levels in the tumor (bars, left axis) were analyzed by qRT-PCR for *SOX10* (d) or for *FGF9* (**e**).

### SMARCA4/SMARCA2 inhibition alters lineage-specific chromatin landscapes

Our results in UM provided a mechanistic understanding of the relationship between BAF and UM-specific TFs. Given the broad activity of FHT-1015 in additional cell types, we hypothesized that BAF ATPase inhibition could have utility across a variety of TF-driven cancer cell types. To test the impact of SMARCA4/SMARCA2 ATPase inhibition on global chromatin accessibility profiles, we performed ATAC-seq across a set of cancer cell lines of diverse origin. As observed in UM, in all cell lines tested, BAF inhibition resulted in a more pronounced loss of accessibility as compared to gain of accessibility (Fig. 5a, Supplementary Fig. 7a). Importantly, FHT-1015 did not impact chromatin accessibility genome-wide, and its effects varied across cell lines (Fig. 5a). For example, the most impacted cell line, the prostate cancer cell line 22RV1, showed a loss of accessibility at 47% of all mapped ATAC-seq peaks,whereby in the colorectal cancer cell line HCT116, only ∼8% of the total ATAC-seq peaks were modulated (Fig. 5a, Supplementary Fig. 7b). Sites of reduced accessibility were preferentially located at promoter-distal rather than promoter-proximal regions (Fig. 5b). The majority of mapped ATAC-seq peaks were unique to each cell line (Supplementary Fig. 7b). Only 7% of all ATAC-seq peaks mapped across all cell lines (10,084 peaks in total) were shared amongst all cell lines, and more than half of these sites were found at promoters (Supplementary Fig. 7b-c). Chromatin regions with reduced accessibility after FHT-1015 treatment were almost entirely cell-line specific (Supplementary Fig. 7d), supporting the notion that BAF plays an important role in maintaining tissue-specific enhancer accessibility^10,37-39^.

**Figure 5:**
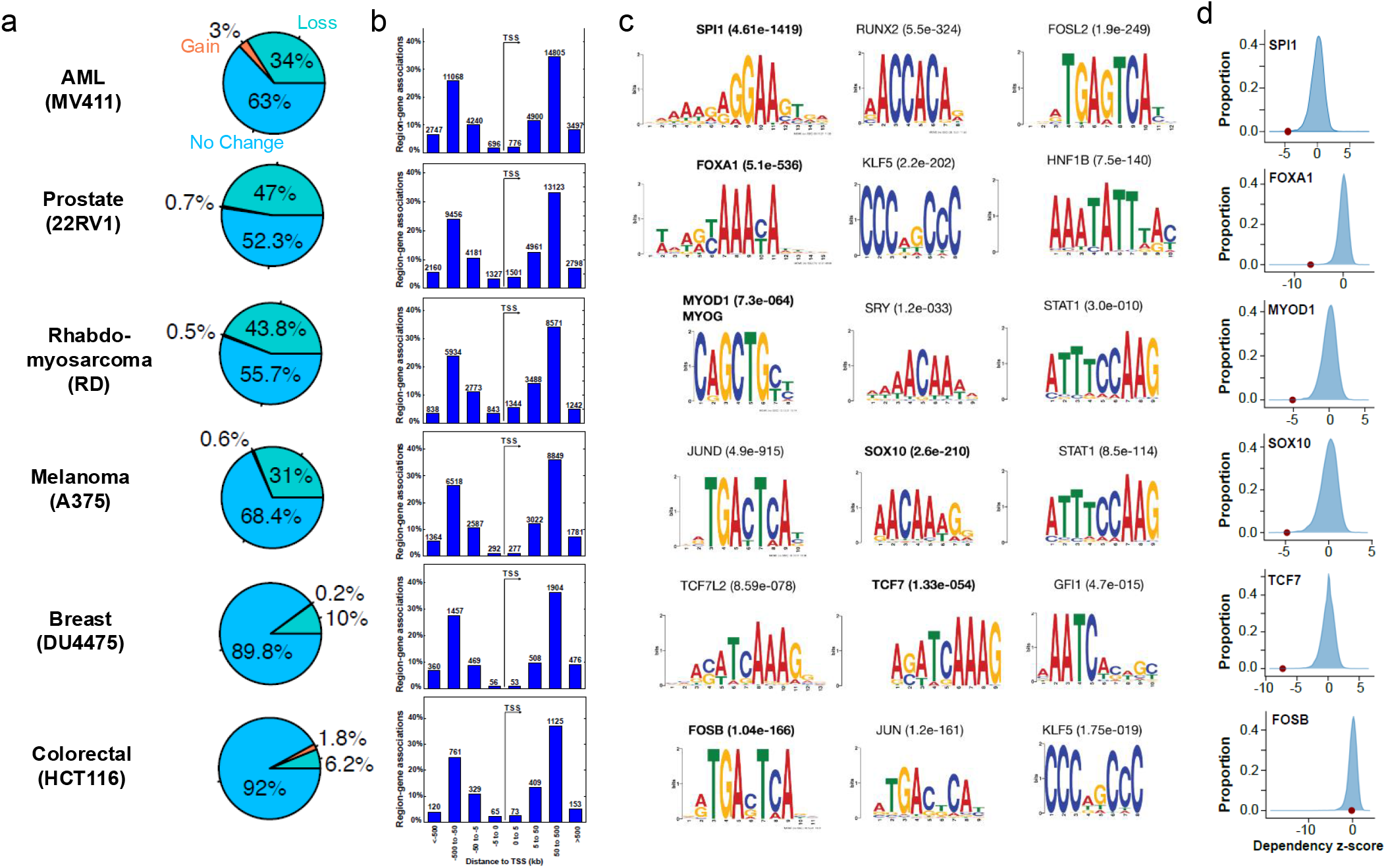
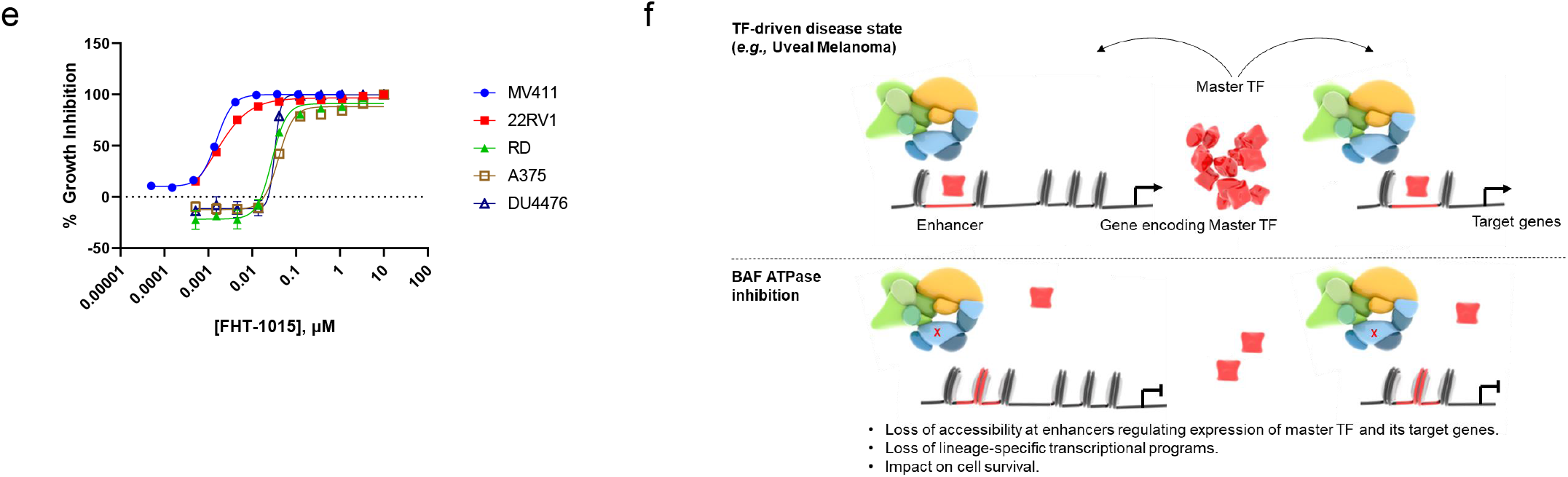
SMARCA4/SMARCA2 ATPase inhibition abrogates accessibility at master TF binding sites. (**a**) Global effects on chromatin accessibility in cancer cell lines treated with FHT-1015 for 4 hours. For each cell line, the percentage of regions with differential accessibility (log_2_FC ≥ 0.5, adjusted p-value < 0.05) is plotted. (**b**) Region-gene association plots showing the distribution of ATAC-seq Cluster 1 (loss of accessibility) peaks in relation to the TSS. For each cell line, Genomic Regions Enrichment of AnnotationsTool (GREAT) algorithm was used to calculate the distance of peaks from the TSS. Number of region-gene associations (0-5 kb, 5-50 kb, 50-500 kb or >500 kb upstream of downstream of the TSS) are found on top of the corresponding bar in the graph. **c**) FHT-1015 reduces chromatin accessibility at lineage transcription factor binding motifs. For each cell line the enrichment of transcription factor binding motif was calculated for regions with loss of chromatin accessibility. The top 3 TF consensus binding motifs most affected and the corresponding enrichment values are indicated. (**d**) For each cell line the CRISPR dependency score from DepMap or Sanger Center studies was z-scored after removal of pan-essential genes. Kernel density estimation plots of all gene dependencies for each cell line are depicted on the right panel with the indicated transcription factor position in the distribution highlighted in red. (**e**) Cells were treated for 7 days with increasing concentrations of FHT-1015 and relative growth was measured by the CTG assay. At day 3, cells were split and fresh media and compound were replenished (n=3; mean ± SD). (**f**) Model summarizing the molecular mechanism of SMARCA4/SMARCA2 ATPase inhibition in TF-driven cancers (*e*.*g*., UM). (Top) in UM cells, BAF ATPase activity maintains chromatin accessibility at enhancers of a master TF, as well as at its target genes, allowing for chromatin occupancy of TFs such as SOX10, MITF, and TFAP2A. This subsequently promotes expression of *SOX10* and downstream transcriptional targets of these transcription factors. (Bottom) Upon BAF ATPase inhibition, chromatin is closed at these important regulatory elements, the chromatin binding of critical TFs is reduced, and gene expression is suppressed.

The BAF complex cooperates with transcription factors to regulate tissue- and disease-specific gene expression programs^14,40^. Consistent with this, FHT-1015 treatment resulted in the closing of chromatin at regions containing transcription factor consensus binding motifs in a manner specific to the cancer cell lineage (Fig. 5c). For example, in the rhabdomyosarcoma cell line, RD, the binding motif for MYOG/MYOD1 transcription factors, critical dependencies for this cell type, was enriched among sites of decreased accessibility (Fig. 5c-d). A similar trend was observed across most cancer cell lineages, whereby the sites of reduced accessibility upon BAFi correlated with binding sites for lineage-specific transcription factors (normally involved in cellular differentiation) with strong cancer dependencies (Fig. 5c-d). Interestingly, one exception was in the colorectal cancer cell line HCT116, the cell line least impacted by FHT-1015, in which no lineage-specific transcription factor motifs were identified at sites of reduced accessibility (Fig. 5c). These results are consistent with the idea that BAF predominantly regulates lineage-specific transcription^14^. In each of the cell lines tested, changes in accessibility ultimately resulted in growth inhibition caused by FHT-1015 (Fig. 5e). Thus, SMARCA4/SMARCA2 ATPase inhibition represents a novel mechanism to alter chromatin accessibility at sites regulated by disease-driving transcription factors. Taken together, these data support a model by which SMARCA4/SMARCA2 inhibition initially results in the loss of lineage-regulated enhancer accessibility,reducing TF occupancy, suppressing tissue-specific transcriptional programs and culminating in growth inhibition of cancer cells (Fig. 5f). This is consistent with several recent publications demonstrating that chromatin accessibility changes occur rapidly, within minutes, upon genetic or pharmacologic inhibition of SMARCA4/SMARCA2, and that the changes in accessibility precede transcriptional effects^39,41,42^.

## Discussion

Here, we describe the discovery and characterization of a novel class of potent and selective SMARCA4/SMARCA2 ATPase inhibitors. With these tool compounds we demonstrate that pharmacologic inhibition of BAF has a selective rather than a genome-wide impact on the chromatin landscape, affecting binding of key transcription factors to their specific enhancers in different cell lineages. While BAF ATPase inhibition has broad activity across a variety of cancer subtypes, uveal melanoma and hematological cancers are exquisitely sensitive at early timepoints. A deeper analysis in uveal melanoma cells showed that SMARCA4/SMARCA2 inhibition results in the loss of enhancer binding of SOX10 and MITF, two master transcriptional regulators in melanocytes. This, in turn, leads to the immediate transcriptional suppression of SOX10 and its downstream gene expression programs, and ultimately apoptosis in UM cells. The example of UM shows that pharmacologic inhibition of SMARCA4/SMARCA2 may provide a therapeutic approach against cancers that depend on lineage transcription factors for survival.

Recently, a distinct series of SMARCA4/SMARCA2 ATPase inhibitors was described^23^. Crystallographic studies demonstrated that these compounds, such as BRM014, bind in the N-terminal RecA lobe of SMARCA2. Multiple lines of evidence presented above (see Fig. 1) strongly suggest that our compound series binds to a different site on SMARCA4/SMARCA2, likely outside of the N-RecA lobe. Despite these differences in the binding mode, the two distinct classes of BAF ATPase inhibitors have similar biological effects, particularly in the inhibition of uveal melanoma cell and tumor growth^43^, underscoring the on-target activity of these compounds, and their utility as chemical probes of BAF ATPase function. Nevertheless, there are some additional differences between our work described here and the study by Rago and colleagues. First, the studies with BRM014 described effects of BAF inhibition only on promoter accessibility, not at distal regulatory elements. While we did observe changes in promoter accessibility, major loss of chromatin accessibility was evident at promoter-distal sites, and in the case of UM cells, these were further characterized as enhancers. BAF complexes occupy both promoters and enhancers to regulate gene expression, but recent work has highlighted the enrichment of canonical BAF complexes at enhancers which is consistent with our observations^6^. As we expect that both series of BAF inhibitors should inhibit all BAF complexes (cBAF, PBAF, and ncBAF), it is possible that the most critical function of BAF is in maintaining enhancer accessibility. Secondly, while Rago and colleagues focused on the role of BAF as an effector of MITF-mediated gene expression, we show here that there are additional important TFs, namely SOX10 and TFAP2A, with which BAF functions to maintain the oncogenic state of UM cells. Functional genomics screens have highlighted the importance of each of these TFs in UM cell viability (Supplementary Fig. 4d), and all three of them have been shown to cooperate with BAF and with each other in melanocyte transcription^33,34^. Finally, we demonstrate here that *SOX10* down-regulation is one of the earliest transcriptional events to occur upon BAFi in UM cell lines, and that overexpression of *SOX10* can override the phenotypic response to BAF inhibition. Intriguingly, Rago and colleagues observed similar effects through the overexpression of *MITF*^43^. How each of these TFs could independently provide resistance to BAFi is currently unclear, although it remains possible that SOX10 and MITF regulate one another^44^, and thus overexpression of one may also lead to overexpression of the other.

While we have detailed the mechanistic response of BAF inhibition in uveal melanoma, there are other disease areas in which this therapeutic modality could be useful. For example, given that SOX10 and MITF are directly affected upon SMARCA4/SMARCA2 inhibition, it is plausible that cutaneous melanomas may respond to these inhibitors. Indeed, chromatin accessibility in cutaneous melanoma cells is also reduced at SOX10 binding sites upon FHT-1015 treatment (Fig. 5). Our cell line screening data suggested that hematologic malignancies and prostate cancer cell lines are also very sensitive to BAFi, and indeed recently published work has demonstrated a very similar mechanism to what is proposed here in regards to SMARCA4/SMARCA2 degradation in prostate cancer^45^. Given the lineage-specific effects of BAF complexes, additional information will be needed to understand the mechanism of action of SMARCA4/SMARCA2 inhibitors in other types of cancers. These studies pave a path to the clinical investigation of BAF ATPase inhibitors for the treatment of UM and other TF-driven cancers.

## Methods

Methods are available in the online version of the paper.

## Data availability

All sequencing data (RNA-seq, PRO-seq, ATAC-seq, ChIP-seq and WES) were deposited at GEO under accession number GEO: GSE241450.

## Acknowledgements

We thank Cigall Kadoch and Craig Peterson for insightful discussions regarding this work. We thank our many Foghorn colleagues for their invaluable help and support. The results discussed here are based, in part, upon data generated by the TCGA Research Network (www.cancer.gov/tcga) and the DepMap project at the Broad Institute (https://doi.org/10.6084/m9.figshare.8061398.v1).

## Author contributions

R.C.C., L.M.M.S., and L.X. designed the study and interpreted the data. R.C.C., K.I., Z.L., L.Z., X.C., G.J.S. and K.F. performed the experiments. L.Z., H.F., J.S., and K.D.B designed and executed the HTS. L.M.M.S., S.T., G.E., J.M., and D.L.L. performed the bioinformatics analyses. R.G.V., N.A., D.H., S.S., K.W., J.V., and D.S.M. designed and synthesized the compounds. R.C.C., A.A., M.H., and L.X. designed the in vivo studies. D.S.M, H.M.C., S.F.B., C.P.D., and R.G.K. provided insight and facilitated study execution. R.C.C., L.M.M.S., S.T. and L.X. wrote the manuscript with insight from co-authors.

