## Supplementary Figures for "Pharmacologic inhibition of BAF chromatin remodeling complexes as a therapeutic approach to transcription factor-dependent cancers"

Supplementary Fig. 1 (Related to Fig. 1). Identification and optimization of a novel class of dual inhibitors of SMARCA4/SMARCA2 ATPase activity.

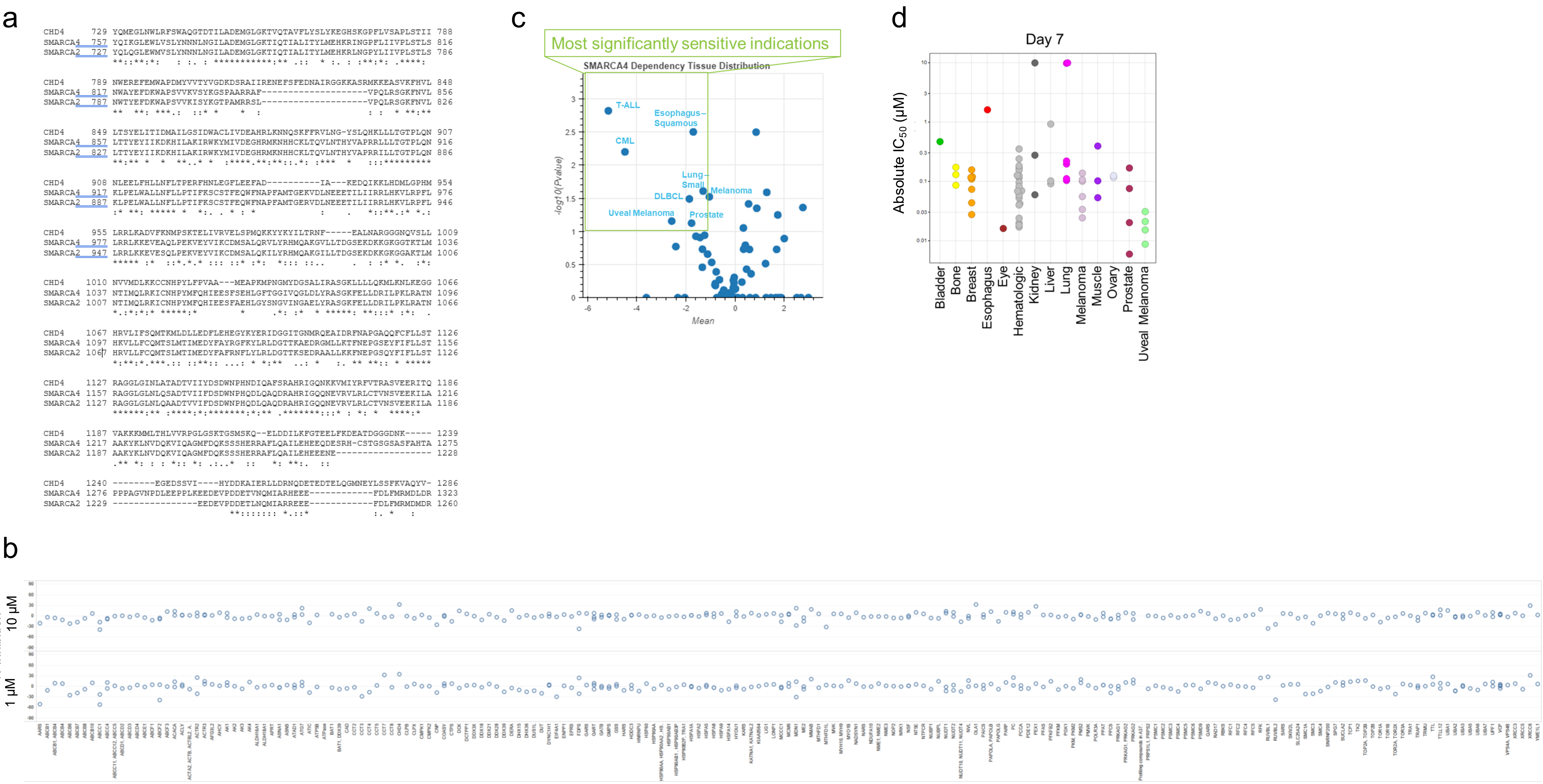

Supplementary Fig. 2 (Related to Fig. 1). The SMARCA4 I1173M mutation confers resistance of MP41 cells to the growth inhibitory effects of FHT-1015.

a

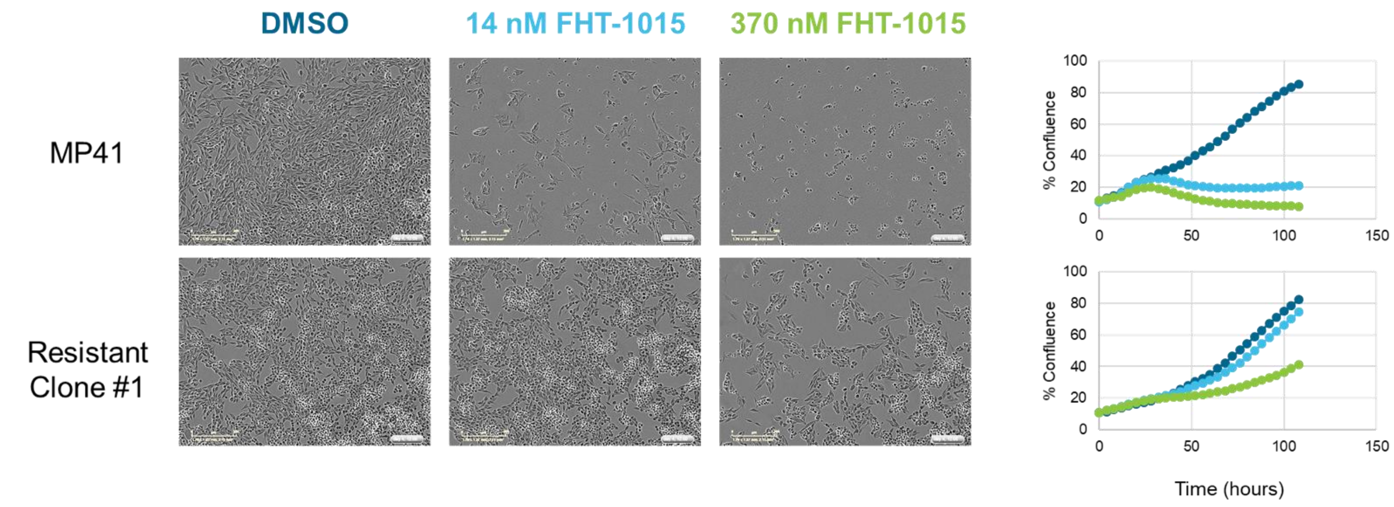


b

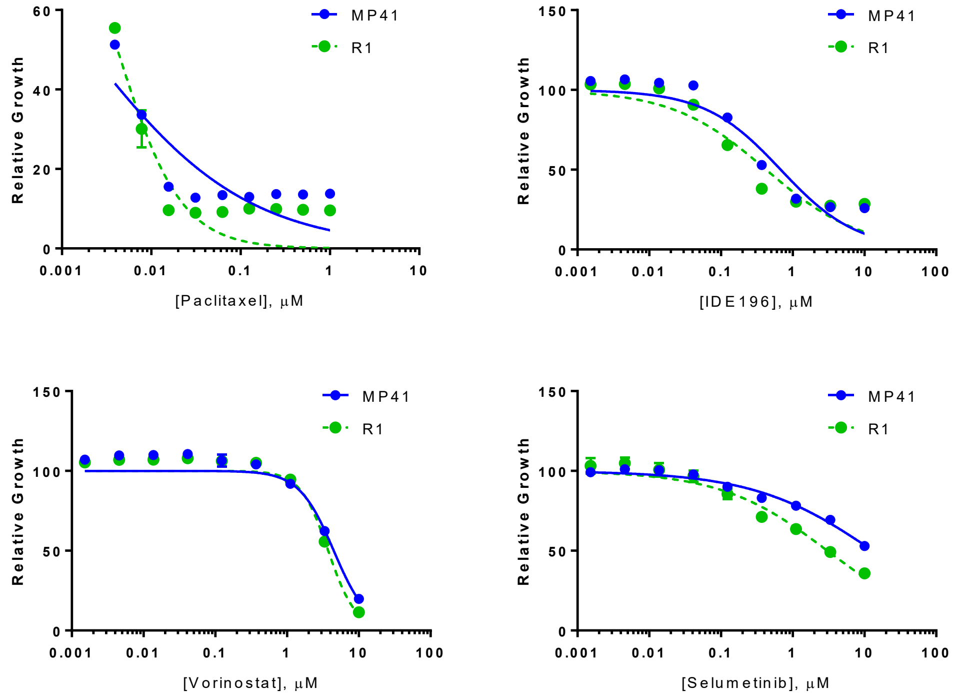


c

| | Gene | Allele Freq. | Mutation | Sample | Other |
|---|---|---|---|---|---|
| Parental MP41 | GNA11 | 0.5 | p.Gln209Leu | MP41 | GoF UM driver |
| | SMARCA4 | 0.5 | p.Arg595Trp | MP41 | unknown |
| | SMARCA4 | 0.5 | p.Thr910Met | MP41 | LoF, ATPase |
| | SMARCA4 | 0.5 | p.Val1469Met | MP41 | unknown, Bromodomain |
| | SMARCA2 | 0.5 | p.Gln236del | MP41 | unknown, polyQ region |

| | Gene | Allele Freq. | Mutation | Sample | Other |
|---|---|---|---|---|---|
| FHT-1015-Resistant Clones | SMARCA4 | 0.5 | p.Ile1173Met | R1 | Corresponds to SMARCA2 I1143 |
| | SMARCA4 | 0.5 | p.Ile1173Met | R2 | Corresponds to SMARCA2 I1143 |
| | SMARCA4 | 0.5 | p.Ile1173Met | R3 | Corresponds to SMARCA2 I1143 |
| | SMARCA4 | 0.5 | p.Ile1173Met | R4 | Corresponds to SMARCA2 I1143 |
| | SMARCA4 | 0.5 | p.Ile1173Met | R5 | Corresponds to SMARCA2 I1143 |
| | SMARCA4 | 0.5 | p.Ile1173Met | R6 | Corresponds to SMARCA2 I1143 |
| | SMARCA4 | 0.5 | p.Ile1173Met | R7 | Corresponds to SMARCA2 I1143 |
| | SMARCA4 | 0.5 | p.Ile1173Met | R8 | Corresponds to SMARCA2 I1143 |
| | SMARCA4 | 0.5 | p.Ile1173Met | R9 | Corresponds to SMARCA2 I1143 |
| | SMARCA4 | 0.5 | p.Gln1387Arg | R7 | |

Supplementary Fig. 3 (Related to Fig. 1). Uveal melanoma cell lines respond to SMARCA4/SMARCA2 ATPase inhibition through an apoptotic mechanism.

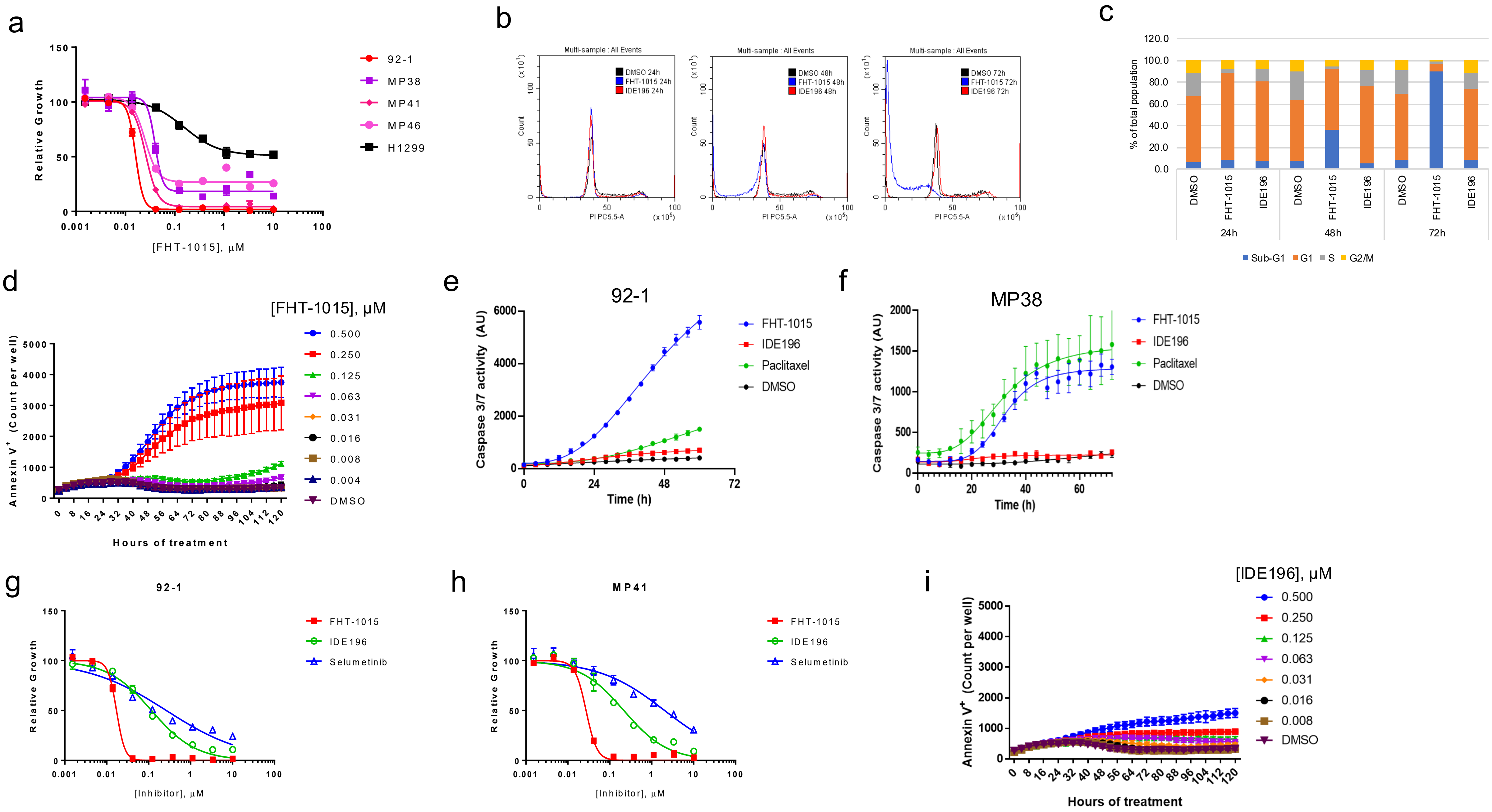

Supplementary Fig. 4 (Related to Fig. 2). BAF ATPase inhibition alters enhancer accessibility of disease-relevant transcription factors.

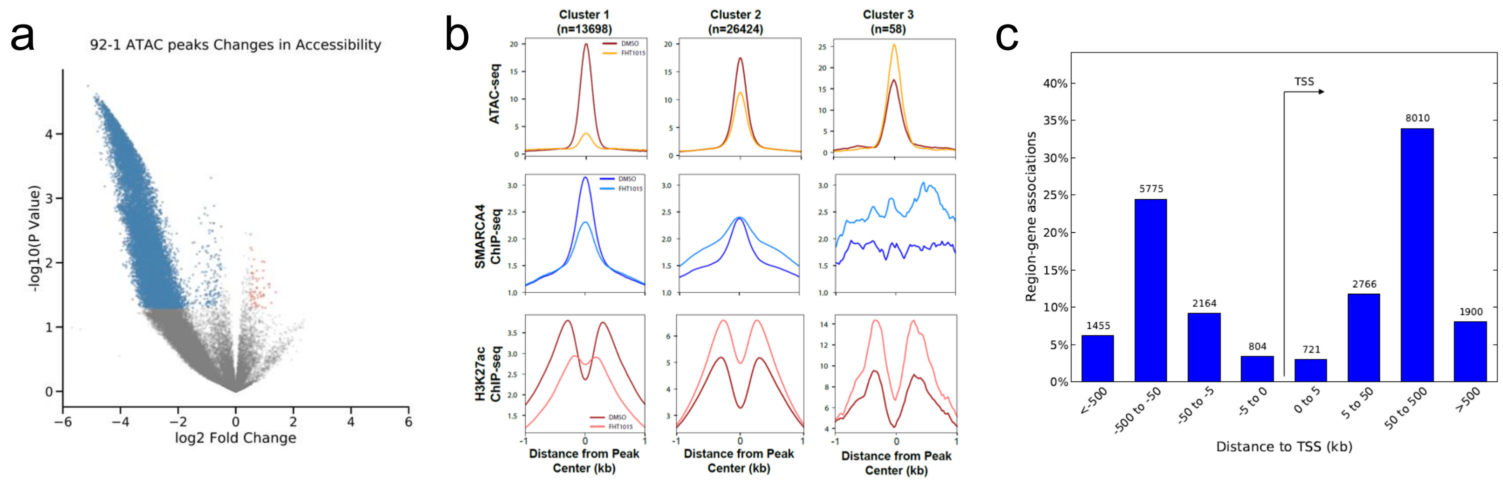


d

### Gene Dependency Rank

| | *GNAQ* | *GNA11* | *SMARCA4* | *SOX10* | *MITF* | *TFAP2A* | *SMARCA2* |
|---|---|---|---|---|---|---|---|
| MEL202 (CERES) | 24 | 2376 | 39 | 2 | 726 | 157 | 15518 |
| 92-1 (DEMETER) | 42 | 15002 | 7 | 72 | 15 | 198 | 13586 |
| OMM1 (DEMETER) | 1580 | 53 | 65 | 17 | 24 | 123 | 12864 |
| MEL285 (DEMETER) | 13044 | 1955 | 3673 | 13718 | 14297 | 15006 | 13842 |

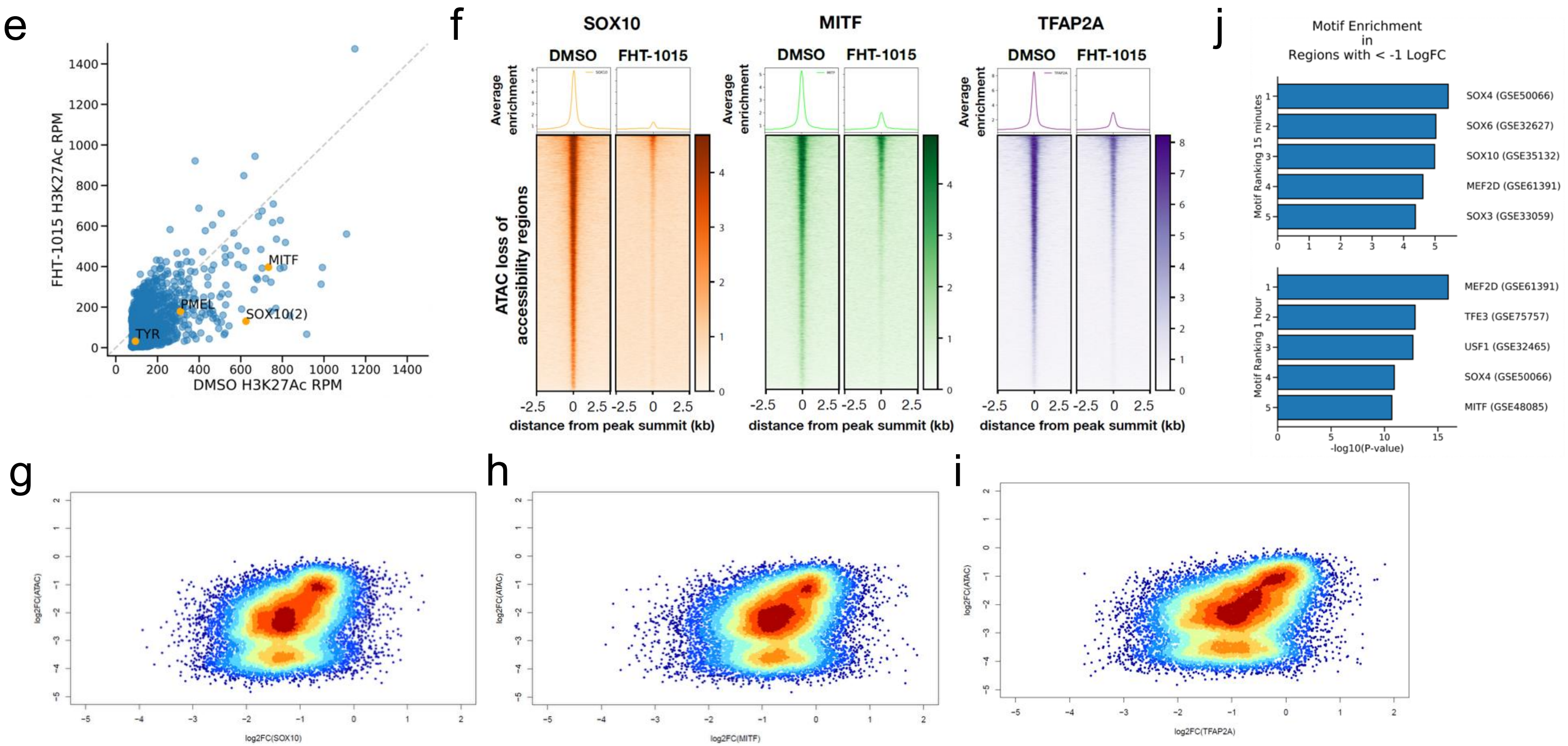

Supplementary Fig. 5 (Related to Fig. 3). BAF ATPase inhibition disrupts the SOX10-MITF transcriptional axis in uveal melanoma.

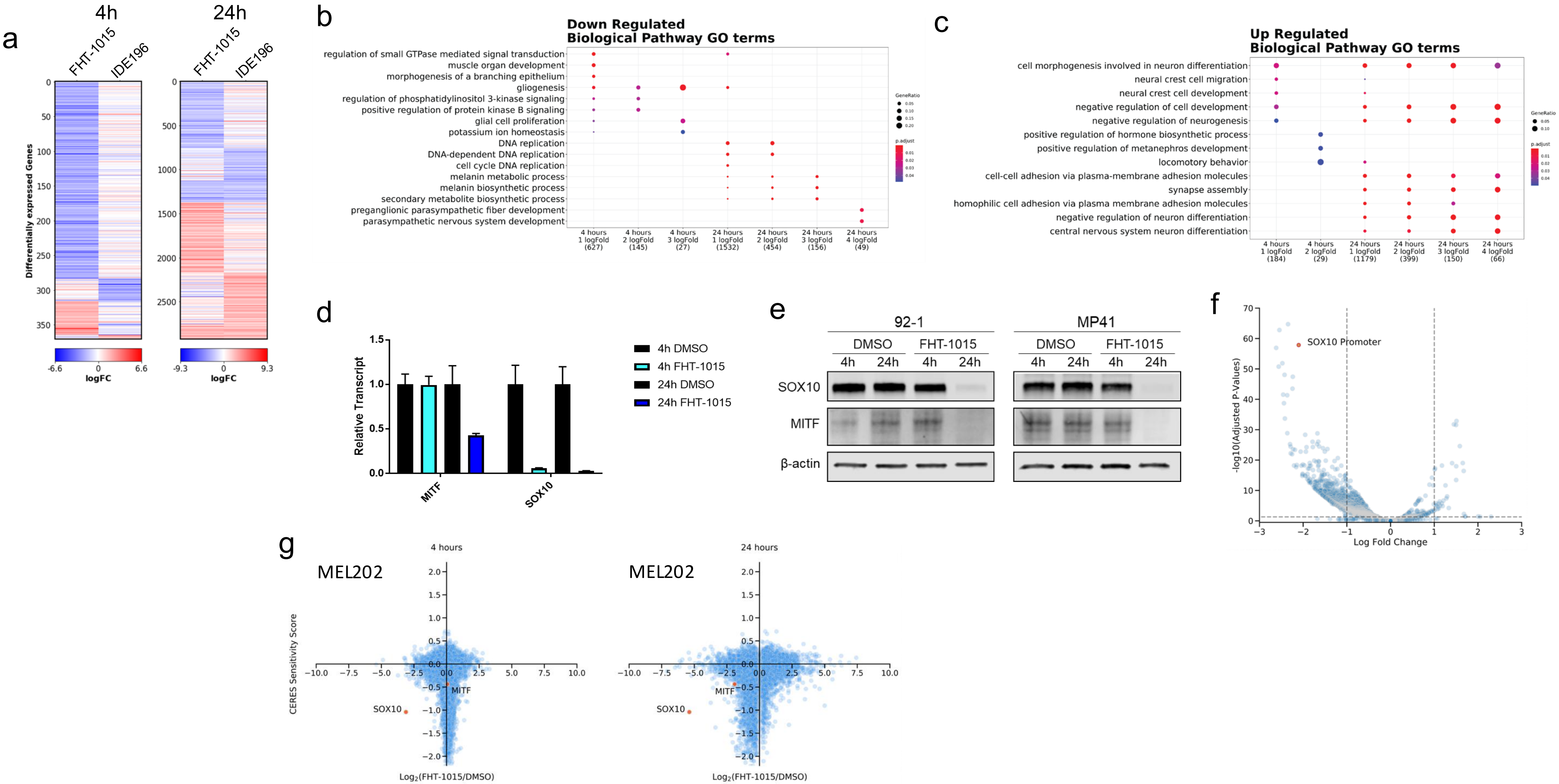

Supplementary Fig. 6 (Related to Fig. 4). BAF ATPase inhibition promotes uveal melanoma tumor regression.

a

| | | |
|---|---|---|
| Physchem Properties | MW | 528.60 |
| | tPSA | 137.21 Å |
| | Log D | 3.36 |
| | pKa | 9.23 |
| Growth Inhibition | 92-1 Cell Proliferation $IC_{50}$ (nM) | 9.61 |
| ADME properties | Kinetic Solubility (µM)<br>Phosphate Buffer/ FaSSIF | 8.8 / 90.8 |
| | Liver Microsome Intrinsic Clearance<br>(Human/ CD-1 Mouse) (µL/min/mg protein) | 17.7/ 17.4 |
| | Plasma Protein Percent Unbound<br>(Human/ CD-1 Mouse) | 0.30/ 0.024 |
| In Vitro Safety | CYP Inhibition $IC_{50}$ (µM)<br>CYP1A2/ CYP2B6/ CYP2C8/ CYP2C9<br>CYP2C19/ CYP2D6/ CYP3A4-M/ CYP3A4-T | >50/ >50/ 15.7/ 7.09<br>39.9/ >50/ >50/ >50 |
| | HERG Inhibition $IC_{50}$ (µM) | 28.1 |
| Mouse PK Parameters | CL (mL/min/kg) | 0.985 |
| | $V_{dss}$ (L/kg) | 0.207 |
| | iv $t_{1/2}$ (h) | 2.09 |
| | po $C_{max}$ (ng/mL) | 25300 |
| | po $AUC_{0\text{-}last}$ (ng•h/mL) | 103977 |
| | Bioavailability %F | ~100% |

b

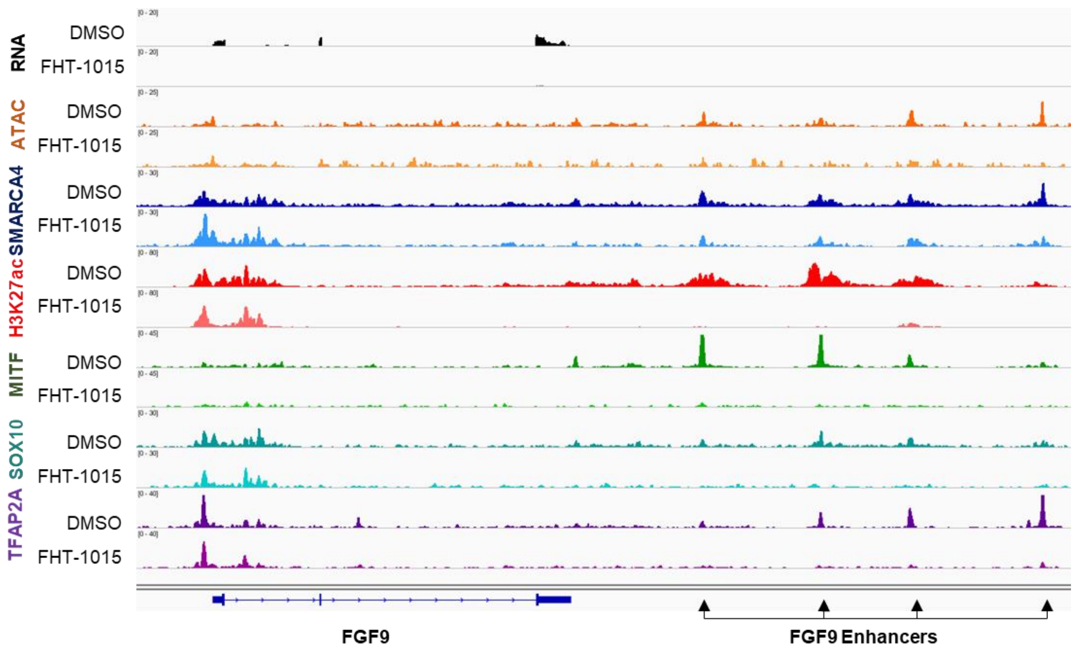

Supplementary Fig. 7 (Related to Fig. 5). SMARCA4/SMARCA2 ATPase inhibition abrogates accessibility at master TF binding sites.

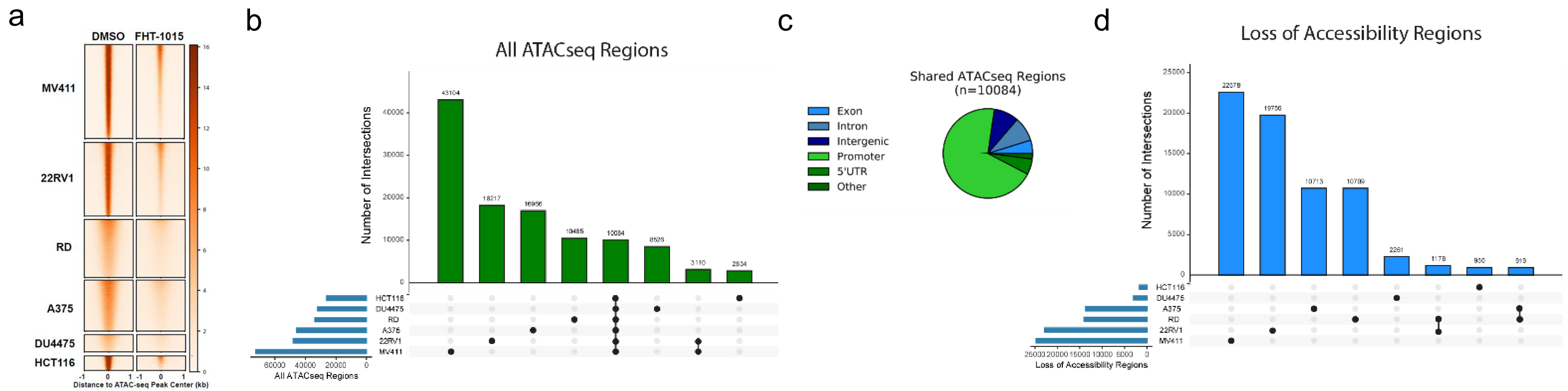
