## Supplementary Text for "Pharmacologic inhibition of BAF chromatin remodeling complexes as a therapeutic approach to transcription factor-dependent cancers"

**SUPPLEMENTARY FIGURE LEGENDS**

**Supplementary Figure 1. Identification and optimization of a novel class of dual inhibitors of SMARCA4/SMARCA2 ATPase activity.** (**a**) Sequence alignment of the ATPase domains of SMARCA4, SMARCA2, and CHD4 demonstrate high similarity between these proteins (SMARCA4 vs. SMARCA2 >90%, SMARCA4/2 vs. CHD4 >50%), rationalizing the use of CHD4 as a related off-target measure of compound selectivity. Protein sequences were retrieved from the Universal Protein Resource website ([www.uniprot.org](http://www.uniprot.org)), and sequences were entered into the Clustal Omega multiple sequence alignment program accessed through The European Bioinformatics Institute website ([www.ebi.ac.uk](http://www.ebi.ac.uk)). (**b**) FHT-2344 was tested against a panel of ATPases in A375 cell lysates using the KiNativ platform (ActivX). Percent inhibition at 1 µM and at 10 µM are plotted. (**c**) Mean Demeter dependency for cancer cell lines of different tissue types on *SMARCA4* (depmap.org) (**d**) Cell lines were treated for 7-days with FHT-1015, and growth relative to DMSO-treatment was measured by CTG. Absolute IC_50_ values are plotted.

**Supplementary Figure 2. The SMARCA4 I1173M mutation confers resistance of MP41 cells to the growth inhibitory effects of FHT-1015.** (**a**) Parental MP41 cells or MP41-derived cells resistant to FHT-1015 were treated with the indicated concentrations of FHT-1015. (Left) Representative micrographs of cells after 120 hours of treatment are shown. (Right) Percent confluence is plotted for MP41 (top) or resistant cells (bottom) at the indicated period of time after treatment with DMSO (dark blue), 14 nM FHT-1015 (light blue), or 370 nM FHT-1015 (green). Confluence was measured in real time on an Incucyte Zoom. (**b**) Parental MP41 or MP41-derived cells that are resistant to FHT-1015 (R1) were treated for 3 days with Paclitaxel, IDE196, Vorinostat, or Selumetinib, and growth relative to a DMSO-control was measured by CTG. (**c**) Notable mutations identified by whole-exome sequencing in parental MP41 cells (top) or MP41-derived isolates that are resistant to growth in the presence of FHT-1015 (bottom). The *SMARCA4* I1173M mutation was identified in all 9 FHT-1015-resistant clones that were sequenced. All mutations highlighted in the table, including T910M in the wild-type MP41, are heterozygous (variant allele frequency is 0.5).

**Supplementary Figure 3. Uveal melanoma cell lines respond to SMARCA4/SMARCA2 ATPase inhibition through an apoptotic mechanism.** (**a**) UM or NSCLC cell lines were treated with a dose titration of FHT-1015 for 3 days, and growth relative to a DMSO-control was measured by CTG. (**b-c**) Histograms (**b**) and quantification (**c**) of cell cycle distributions of cells treated for 1, 2, or 3 days with DMSO, 100 nM FHT-1015, or 1 µM IDE196. (**d**) 92-1 cells were treated with DMSO or a dose range of FHT-1015, and Annexin V positivity was measured in real time using an Incucyte Zoom. (**e-f**) 92-1 (**e**) or MP38 (**f**) UM cells were treated with 111 nM FHT-1015, IDE196, Paclitaxel, or DMSO, and Caspase 3/7 activity was measured in real time using an Incucyte Zoom. (**g-h**) 92-1 (**g**) or MP41 (**h**) UM cells were treated for 3 days with dose titrations of FHT-1015, IDE196, or Selumetinib, and growth relative to a DMSO-control was measured by CTG. (**i**) 92-1 cells were incubated with the indicated dose of IDE196, and Annexin V positivity was measured in real time using an Incucyte Zoom.

**Supplementary Figure 4. BAF ATPase inhibition alters enhancer accessibility of disease-relevant transcription factors.** (**a**) Differential accessibility of ATAC-seq peaks in 92-1 UM cells after treatment with 100 nM FHT-1015 for 4 hours. Coverage values for regions identified consistently among replicates of ATAC-seq experiment were retrieved, and differential changes in accessibility were calculated corresponding consistency. Blue dots indicate regions with significantly reduced accessibility (log_2_ fold change < 0.5 (p < 0.05); red dots indicate regions with significantly increased accessibility (log_2_ fold change > 0.5 (p < 0.05). (**b**) 92-1 ATAC-seq peaks were classified as loss of accessibility (Cluster 1), not changing (Cluster 2) and gain of accessibility (Cluster 3). Average signal for ATAC-seq, SMARCA4 ChIP-seq and H3K27Ac ChIP-seq was calculated and depicted as anchor plot centered at the ATAC-seq peak maximum value (flanking 1 kb upstream or downstream). (**c**) Region-gene association plots showing the distribution of ATAC-seq Cluster 1 (loss of accessibility) peaks in relation to the TSS. For 92-1 ATAC-seq, Genomic Regions Enrichment of Annotations Tool (GREAT) algorithm was used to calculate the distance of Cluster 1 peaks from the TSS. Number of region-gene associations (0-5 kb, 5-50 kb, 50-500 kb or >500 kb upstream of downstream of the TSS) are found on top of the corresponding bar in the graph. (**d**) For each cell line, all genes were ranked by their CERES or DEMETER score (depmap.org), and pan-essential genes were filtered out. The ranked gene dependency is listed for the given genes in the table. Green highlighting represents genes within the top 750 strongest dependencies. All other gene ranks are highlighted in red. (**e**) Loss of H3K27ac at individual enhancers regulating melanocyte biology (highlighted in orange). Enhancers in 92-1 cells were identified using ROSE with H3K27ac ChIP-seq data. For each enhancer identified, the corresponding gene was assigned based on proximity and the normalized levels of H3K27Ac for DMSO (x-axis) or FHT-1015 treated cells (y-axis) are plotted. (**f**) Average ChIP-seq enrichment of SOX10, MITF, and TFAP2A at Cluster 1 loss of accessibility sites following treatment of 92-1 cells with DMSO or 100 nM FHT-1015 for 4 hours. Anchor plots are centered on ChIP-seq peaks for each indicated TF. (**g-i**) Correlation of differential chromatin accessibility (measured by ATAC-seq) with differential occupancy of SOX10 (**g**), MITF (**h**), or TFAP2A (**i**) (measured by ChIP-seq) in response to a 4 hour treatment with 100 nM FHT-1015. (**j**) Motif enrichment in downregulated bidirectional transcription regions in MP46. Motifs enriched and corresponding p-values in regions identified through PRO-seq showing at least 2-fold decrease in signal at the indicated timepoints are indicated, motif information is provided on the left.

**Supplementary Figure 5. BAF ATPase inhibition disrupts the SOX10-MITF transcriptional axis in uveal melanoma.** (**a**) Differential gene expression of 92-1 cells treated with 100 nM FHT-1015 or 1 µM IDE196 after 4 and 24 hours. Genes modulated at least 2-fold with adjusted p-value < 0.05 in any condition were used. (**b-c**) Gene ontology (GO term) overrepresentation analysis for down- (**b**) or up- (**c**) regulated genes in 92-1 treated with FHT-1015. For each time of treatment (4 or 24 hours) the genes showing statistically significant down- (**b**) or up- (**c**) regulation stronger than indicated in the bottom annotation were identified. GO term overrepresentation analysis was performed for each group of genes using Biological Pathway GO terms. The top overrepresented terms for each group of genes is indicated together with number of genes identified and corresponding statistical significance. (**d**) 92-1 cells were treated for 4h or 24h with FHT-1015 and relative expression of *SOX10* or *MITF* was measured by qRT-PCR. (**e**) 92-1 or MP41 cells were treated as in (d), and SOX10 or MITF protein expression was measured by Western blot. (**f**) MP46 cells were treated for 1 hour with 100 nM FHT-1015 and nascent transcripts were measured by PRO-seq. Differentially expressed nascent transcripts are plotted. *SOX10* is highlighted a significantly reduced nascent transcript. (**g**) MEL202 UM cells were treated with DMSO or 100 nM FHT-1015 for 4h or 24h and gene expression changes were plotted against the DEMETER dependency score (depmap.org). *SOX10* is highlighted as a strong dependency with rapid and robust suppression of gene expression upon FHT-1015 treatment.

**Supplementary Figure 6. BAF ATPase inhibition promotes uveal melanoma tumor regression.** (**a**) Table of ADME properties for FHT-2344. (**b**) Genome browser view of the *FGF9* locus showing enrichment of accessible chromatin, SMARCA4, MITF, SOX10, and TFAP2A at broad H3K27ac peaks, and regulation of these factors by FHT-1015.

**Supplementary Figure 7. SMARCA4/SMARCA2 ATPase inhibition abrogates accessibility at master TF binding sites.** (**a**) Changes in chromatin accessibility upon FHT-1015 treatment. For each cell line indicated on the left, chromatin accessibility heatmaps with normalized coverage ATAC-seq signal tracks centered at the called peak are shown for regions losing accessibility, corresponding regions are represented for DMSO and FHT-1015 treated experiments. (**b**) The majority of chromatin accessible regions identified by ATAC-seq are cell line specific. Upset plot indicating the overlap in open chromatin regions between all cell lines indicated, approximately 10000 regions are shared between all cell lines. (**c**) Shared chromatin accessible regions are enriched for promoter proximal regions. Shared regions identified in (c) were annotated according to genomic context and proportions are represented as a pie chart. (**d**) Chromatin accessible regions identified by ATAC-seq as losing chromatin accessibility are almost exclusively cell line specific. Upset plot indicating the overlap in open chromatin regions that loose accessibility upon treatment with FHT-1015 for 4 hours between all cell lines indicated, no regions are shared among all cell lines.

**ONLINE METHODS**

**Protein constructs**

Full-length SMARCA2 (1-1590, C-terminal 6-His FLAG tag) and SMARCA4 (1-1647, C-terminal 6-His FLAG tag) and CHD4(1-1912, N-terminal FLAG and C-terminal-6-His tag) were produced from Viva Biotech. SMARCA2 I1143M (FL, 1-1590, C-terminal 6-His FLAG tag) and SMARCA4 I1173M (FL, 1-1647, C-terminal His FLAG tag) were constructed and produced at Viva Biotech.

**HTS**

The ATPase catalytic activity of SMARCA2 was measured using the ADP-Glo™ detection reagent (Promega cat# V9102). The assay reaction mixture (10µl) contained 20nM SMARCA2, 10nM salmon sperm DNA (UltraPure™ Salmon Sperm DNA Solution, Invitrogen cat# 15632011), and 400µM ATP in the ATPase assay buffer, which consisted of 20mM Tris, pH8, 20mM MgCl2, 50mM NaCl, 0.1% Tween-20, and 1mM fresh DTT (Pierce™ DTT (Dithiothreitol), ThermoFisher Scientific cat# 20290). The final DMSO concentration was 1% v/v.

The reaction is initiated by the addition of 2µl ATP/DNA solution to a 3µl ATPase solution on low volume white Proxiplate-384 plus plate (PerkinElmer cat# 6008280) which were prepared to achieve a compound dose of 10 µl. The plates are then incubated at room temperature for 1hr. Next 5µl of ADP-Glo™ Reagent is added into the reaction and incubated at room temperature for 40 minutes. Then, 10µl of Kinase Detection Reagent is added to convert ADP to ATP and incubated at room temperature for 60 minutes followed by luminescence measurement with an Envision plate-reader.

Approximately 165,000 compounds were screened. The 0% inhibition effect was established using no compound in 32 wells per plate. The 100% inhibition effect was established using no SMARCA2 enzyme in 32 wells per plate. A hit criteria was applied using three times the standard deviation plus the mean of the percent effect for all compound wells. Thus, 497 compounds demonstrated >26.5% inhibition for a hit rate of 0.30%. A cohort of approximately 500 compounds were retested at 10 uM in triplicate using the same assay format as described for the primary HTS. A confirmation rate of 42% was achieved using an activity cutoff >26%. Correlation between the triplicates was excellent. Subsequently, 211 compounds were selected for IC50 evaluation.

**SMARCA2 and SMARCA4 ADP-glo assay for IC50 evaluation**

The ATPase catalytic activity of SMARCA4 or SMARCA2 was measured using the ADP-Glo™ detection reagent (Promega cat# V9102). The assay reaction mixture (5µl) contained 5nM SMARCA2 or SMARCA4, 10nM salmon sperm DNA (UltraPure™ Salmon Sperm DNA Solution, Invitrogen cat# 15632011), and 400µM ATP in the ATPase assay buffer, which consisted of 20mM Tris, pH8, 20mM MgCl2, 50mM NaCl, 0.1% Tween-20, and 1mM fresh DTT (Pierce™ DTT (Dithiothreitol), ThermoFisher Scientific cat# 20290).

The reaction is initiated by the addition of 2µl ATP/DNA solution to a 3µl ATPase solution on low volume white Proxiplate-384 plus plate (PerkinElmer cat# 6008280) which are prepared with the appropriate amount of compound dose. The plates are then incubated at room temperature for 2hr. Next 5µl of ADP-Glo™ Reagent is added into the reaction and incubated at room temperature for 40 minutes. Then, 10µl of Kinase Detection Reagent is added to convert ADP to ATP and incubated at room temperature for 60 minutes followed by luminescence measurement with an Envision plate-reader.

**CHD4 ADP-glo assay for IC50 evaluation**

The ATPase catalytic activity of CHD4 was measured using the ADP-Glo™ detection reagent. The assay reaction mixture (5µl) contained 25nM CHD4, 100nM salmon sperm DNA, and 400µM ATP in the ATPase assay buffer, which consisted of 20mM Tris, pH8, 20mM MgCl2, 50mM NaCl, 0.1% Tween-20, and 1mM fresh DTT, as above.

The reaction is initiated by the addition of 2µl ATP/DNA solution to a 3µl ATPase solution on low volume white Proxiplate-384 plus plate (PerkinElmer cat# 6008280) which are prepared with the appropriate amount of compound dose. The plates are then incubated at room temperature for 1hr. Next 5µl of ADP-Glo™ Reagent is added into the reaction and incubated at room temperature for 40 minutes. Then, 10µl of Kinase Detection Reagent is added to convert ADP to ATP and incubated at room temperature for 60 minutes followed by luminescence measurement with an Envision plate-reader.

**PK/LDH Coupled Enzyme Assay**

Enzymes and reaction mix were prepared in the assay buffer (20mM Tris HCl buffer at pH 8.0 containing 20mM MgSO4, 50mM KCl, 0.01% Tween-20, and 1mM DTT). Each assay well (50.5µl) contained 0.5µl of variable concentrations of inhibitors or DMSO, and 5nM enzyme together with 200nM salmon-sperm DNA and a reaction mix consisting of final concentrations of 1 mM ATP substrate, 2.5 mM PEP, 10 U/ml PK/LDH and 60 µM NADH. The reaction was performed in Corning black 384-well plates at 23°C and was initiated by adding reaction mix to the assay plate. Fluorescence values at Ex 345nm Em 476nm were measured with the Envision plate reader at 10 min time intervals over the course of 3 hours. The linear rate of NADH loss was determined at each concentration of inhibitor and thus percent inhibition calculated using the rate of NADH loss of the plate controls. For this kinetic read assay the zero percent inhibition controls consist of no compound (1% DMSO) and the one hundred percent inhibition controls consist of no enzyme.

**Medicinal Chemistry**

**Preparation of 3-(dimethylsulfamoyl)-*N*-[2-oxo-2-[(4-phenylthiazol-2-yl)amino]ethyl]benzamide (FHT-185, Compound 1)**

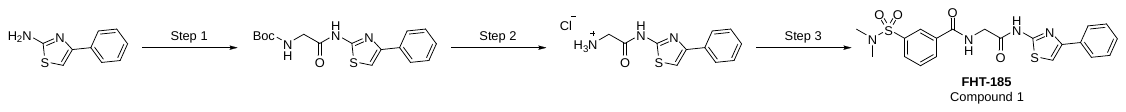

*Step 1: Preparation of tert-butyl N-[2-oxo-2-[(4-phenylthiazol-2-yl)amino]ethyl]carbamate*

To a solution of 2-(tert-butoxycarbonylamino)acetic acid (1.00 g, 5.71 mmol) in DMF (20 mL) was added 1-[bis(dimethylamino)methylene]-1H-1,2,3-triazolo[4,5-b]pyridinium-3-oxide hexafluorophosphate (2.39 g, 6.28 mmol) and *N,N*-diisopropylethylamine (3.0 mL, 17.1 mmol). After stirring for 10 minutes, 4-phenylthiazol-2-amine (1.01 g, 5.71 mmol) was added to the reaction mixture. After stirring at 30 °C for 12 h, the reaction solution was poured over water and extracted with ethyl acetate. The organic layer was washed five times with aqueous citric acid, one with brine, dried over Na_2_SO_4_, and concentrated under reduced pressured. The residue was purified via silica gel flash chromatography to afford *tert*-butyl *N*-[2-oxo-2-[(4-phenylthiazol-2-yl)amino]ethyl]carbamate (2.85 g, 49.9% yield) as a yellow solid. ^1^H NMR (400 MHz, CDCl_3_) d 10.08 (brs, 1H), 7.83–7.81 (m, 2H), 7.45–7.41 (m, 2H), 7.36–7.34 (m, 1H), 7.28 (s, 1H), 7.17 (s, 1H), 4.04 (s, 2H), 1.51 (s, 9H). LCMS (ESI) m/z: = 333.9 (M + H^+^).

*Step 2: Preparation of 2-amino-N-(4-phenylthiazol-2-yl)acetamide hydrochloride salt*

To a solution of *tert*-butyl *N*-[2-oxo-2-[(4-phenylthiazol-2-yl)amino]ethyl]carbamate (0.100 g, 0.285 mmol) in 1,4-dioxane (5 mL) was added a solution of HCl (4 M in 1,4-dioxane, 5.00 mL, 20 mmol). After stirring at 30 °C for 1h, the reaction mixture was concentrated under reduced pressured to afford 2-amino-N-(4-phenylthiazol-2-yl)acetamide hydrochloride salt as a white solid, which was utilized in the next step without further purification. LCMS (ESI) m/z: = 233.9 (M + H^+^).

*Step 3: Preparation of 3-(dimethylsulfamoyl)-N-[2-oxo-2-[(4-phenylthiazol-2-yl)amino]ethyl]benzamide (FHT-185, Compound 1)*

To a solution of 3-(dimethylsulfamoyl)benzoic acid (0.076 g, 0.330 mmol) in dichloromethane (10 mL) was added 1-[bis(dimethylamino)methylene]-1H-1,2,3-triazolo[4,5-b]pyridinium-3-oxide hexafluorophosphate (0.126 mg, 0.330 mmol) and *N,N*-diisopropylethylamine (0.21 mL, 1.2 mmol). After stirring for 10 minutes, 2-amino-*N*-(4-phenylthiazol-2-yl)acetamide hydrochloride salt (0.081 g, 0.300 mmol) was added to the reaction mixture. After stirring at 30 °C for 12 h, the reaction solution was poured over water and extracted with ethyl acetate. The organic layer was washed with brine, dried over anhydrous Na_2_SO_4_, and concentrated under reduced pressure. To the resulting residue was added MeOH (20 mL). After stirring for 20 minutes, the mixture was filtered and the solids were dried to afford 3-(dimethylsulfamoyl)-*N*-[2-oxo-2-[(4-phenylthiazol-2-yl)amino]ethyl]benzamide (67 mg, 50.2% yield) as a white solid (FHT-185, Compound 1). ^1^H NMR (400 MHz, DMSO-d_6_) d 12.49 (s, 1H), 9.32 (t, J = 5.6 Hz, 1H), 8.27–8.25 (m, 2H), 7.96–7.90 (m, 3H), 7.81–7.80 (m, 1H), 7.64 (s, 1H), 7.46–7.42 (m, 2H), 7.33–7.30 (m, 1H), 4.23 (d, J = 5.6 Hz, 2H), 2.65 (s, 6H)**.** LCMS (ESI) m/z: = 444.9 (M + H^+^).

**Preparation of 1-methylsulfonylpyrrole-3-carboxylic acid**

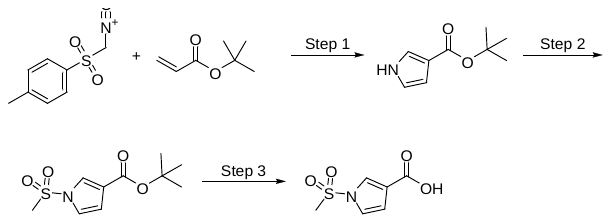

*Step 1. Preparation of tert-butyl 1H-pyrrole-3-carboxylate*

To a mixture of tert-butyl-prop-2-enoate (78.6 mL, 542 mmol) and 1-(isocyanomethylsulfonyl)-4-methylbenzene (106 g, 542 mmol) in THF (1300 mL) was added 60% NaH in mineral oil (25.97 g, 649 mmol) slowly at 30 °C over 1 hour and then heated to 70 °C. After 2 h, the reaction mixture was poured into saturated aqueous NH_4_Cl solution and extracted three times with ethyl acetate. The combined organic phase was washed twice with brine, dried with anhydrous Na_2_SO_4_, filtered, and concentrated under reduced pressure to afford a residue. The residue was purified by silica gel flash chromatography to afford tert-butyl 1H-pyrrole-3-carboxylate (41.5 g, 43% yield) as a yellow solid. ^1^H NMR (400 MHz, CDCl_3_) δ 8.36 (br s, 1H), 7.35 - 7.25 (m, 1H), 6.71 - 6.62 (m, 1H), 6.59 - 6.49 (m, 1H), 1.48 (s, 9H). LCMS (ESI) m/z: = 180.4 (M + Na^+^).

*Step 2. Preparation of tert-butyl 1-(methylsulfonyl)-1H-pyrrole-3-carboxylate*

To a cooled solution (0 °C) of tert-butyl 1H-pyrrole-3-carboxylate (40.5 g, 242 mmol) in THF (1500 mL) was added a 1 M solution of NaHMDS (484 mL, 484 mmol). After stirring at 0 °C for 30 min, methanesulfonyl chloride (28.1 mL, 363 mmol) was slowly added and the mixture was warmed to 30 °C. After 16 h, the reaction mixture was slowly poured into saturated aqueous NH_4_Cl solution and extracted three times with ethyl acetate. The combined organic layers were washed twice with brine, dried with anhydrous Na_2_SO_4_, filtered, and concentrated under reduced pressure to afford a residue. The residue was purified by silica gel chromatography to afford a yellow solid. The yellow solid was triturated with methyl tert-butyl ether at room temperature, stirred for 20 minutes, filtered, and dried in vacuum to afford tert-butyl 1-(methylsulfonyl)-1H-pyrrole-3-carboxylate (25.7 g, 43 % yield) as a white solid. ^1^H NMR (400 MHz, CDCl_3_) δ 7.66-7.64 (m, 1H), 7.10 - 7.08 (m, 1H), 6.73-6.71 (m, 1H), 3.21 (s, 3H), 1.56 (s, 9H).

*Step 3. Preparation of 1-methylsulfonylpyrrole-3-carboxylic acid*

To a mixture of tert-butyl 1-(methylsulfonyl)-1H-pyrrole-3-carboxylate (25.7 g, 105 mmol) in 1,4-dioxane (100 mL) was added a 4 M solution of HCl in 1,4-dioxane (400 mL, 1.6 mol) at 15 °C. After stirring at 15 °C for 14 h, the reaction mixture was concentrated under reduced pressure to afford a residue. The residue was triturated with methyl tert-butyl ether at 15 ^o^C for 16 h. The mixture was filtered and dried in vacuum to afford 1-(methylsulfonyl)-1H-pyrrole-3-carboxylic acid (18.7 g, 94 % yield) as a white solid. ^1^H NMR (400 MHz, Methanol-d_4_) δ 7.78 - 7.77 (m, 1H), 7.25 - 7.23 (m, 1H), 6.72 - 6.70 (m, 1H), 3.37 (s, 3H). LCMS (ESI) m/z: = 189.8 (M + H^+^).

**Preparation of (*S*)-4-amino-*N*-(4-(methylthio)-1-oxo-1-((4-(3-(pyridin-4-yl)phenyl)thiazol-2-yl)amino)butan-2-yl)benzamide (FHT-2344, Compound 2)**

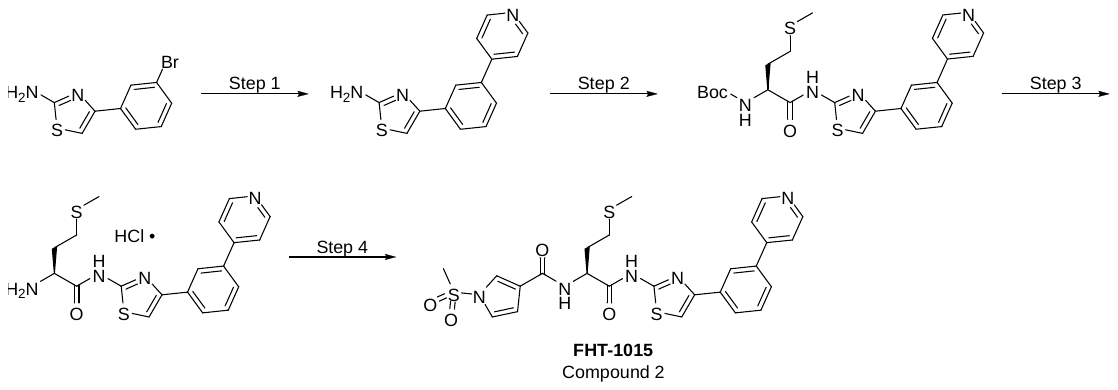

*Step 1: Preparation of 4-[3-(4-pyridyl)phenyl]thiazol-2-amine*

4-(3-bromophenyl)thiazol-2-amine (20.0 g, 78.4 mmol), 4-pyridylboronic acid (28.9 g, 239 mmol), Pd(DPPF)Cl_2_ (2.56 g, 3.92 mmol) and K_3_PO_4_ (66.6 g, 314 mmol) were suspended in a mixture of 1,4-dioxane (240 mL) and H_2_O (24 mL). After stirring at 80 ⁰C for 7 h, the mixture was cooled to ambient temperatures and diluted with water. The aqueous phase was extracted three times with ethyl acetate. The combined organic layers were washed with brine, dried over anhydrous Na_2_SO_4_, filtered and concentrated under reduced pressure. The residue was triturated using a mixture of dichloromethane (30 mL) and MTBE (100 mL) and stirred for 5 minutes. The resulting precipitates were collected through filtration and washed with additional MTBE to afford 4-[3-(4-pyridyl)phenyl]thiazol-2-amine (16.2 g, 78.0% yield) as a yellow solid. LCMS (ESI) m/z: = 254.0 (M + H^+^).

*Step 2: Preparation of* *tert-butyl N-[(1S)-3-methylsulfanyl-1-[[4-[3-(4-pyridyl)phenyl]thiazol-2-yl]carbamoyl]propyl]carbamate*

To a mixture of 4-[3-(4-pyridyl)phenyl]thiazol-2-amine (12.6 g, 49.7 mmol) and (2*S*)-2-(*tert*-butoxycarbonylamino)-4-methylsulfanyl-butanoic acid (18.6 g, 74.6 mmol) in dichloromethane (900 mL) was added 2-ethoxy-1-ethoxycarbonyl-1,2-dihydroquinoline (24.6 g, 99.5 mmol). After stirring at RT for 2 h, the reaction mixture was concentrated under reduced pressure. The residue was triturated with dichloromethane (100 mL) followed by addition of MeOH (200 mL) to afford *tert*-butyl *N*-[(1*S*)-3-methylsulfanyl-1-[[4-[3-(4-pyridyl)phenyl]thiazol-2-yl]carbamoyl]propyl]carbamate (11.7 g, 47.7% yield) as a white solid. ^1^H NMR (400 MHz, DMSO-d_6_) d 12.39 (s, 1H), 8.68–8.66 (m, 2H), 8.30 (s, 1H), 8.02–7.99 (m, 1H), 7.83 (s, 1H), 7.76–7.74 (m, 3H), 7.61–7.57 (m, 1H), 7.28 (d, J = 7.6 Hz, 1H), 4.31–4.30 (m, 1H), 2.65–2.44 (m, 2 H), 2.06 (s, 3H) 2.01–1.85 (m, 2H), 1.38 (s, 9H). LCMS (ESI) m/z: = 485.1 (M + H^+^).

*Step 3: Preparation of (S)-2-amino-4-(methylthio)-N-(4-(3-(pyridin-4-yl)phenyl)thiazol-2-yl)butanamide hydrochloride*

To a solution of *tert*-butyl *N*-[(1*S*)-3-methylsulfanyl-1-[[4-[3-(4-pyridyl)phenyl]thiazol-2-yl]carbamoyl]propyl]carbamate (11.5 g, 23.7 mmol) in MeOH (50 mL) was slowly added a solution of HCl (4 M in 1,4-dioxane, 100 mL, 400 mmol). After stirring at RT for 1 h, the reaction mixture was poured into MTBE. The resulting precipitates were collected by vacuum filtration to afford (S)-2-amino-4-(methylthio)-N-(4-(3-(pyridin-4-yl)phenyl)thiazol-2-yl)butanamide hydrochloride (9.99 g, quant. yield) as a yellow solid. LCMS (ESI) m/z: = 385.0 (M + H^+^).

*Step 4: Preparation of (S)-1-(methylsulfonyl)-N-(4-(methylthio)-1-oxo-1-((4-(3-(pyridin-4-yl)phenyl)thiazol-2-yl)amino)butan-2-yl)-1H-pyrrole-3-carboxamide (FHT-1015, Compound 2)*

To a solution of (S)-2-amino-4-(methylthio)-N-(4-(3-(pyridin-4-yl)phenyl)thiazol-2-yl)butanamide hydrochloride (2.00 g, 4.75 mmol) and 1-methylsulfonylpyrrole-3-carboxylic acid (0.899 g, 4.75 mmol) in DMF (20 mL) was added *N*-(3-dimethylaminopropyl)-*N*′-ethylcarbodiimide hydrochloride (1.37 g, 7.13 mmol), 1-hydroxybenzotriazole (0.963 g, 7.13 mmol) and *N*,*N*-diisopropylethylamine (3.31 mL, 19.0 mmol). After stirring at RT for 3 h, the reaction mixture was poured over H_2_O. The resulting precipitates were collected via vacuum filtration. The solid was initially triturated with MeOH (20 mL) and the precipitate was collected via vacuum filtration. The resulting solid were crystallized from MeOH (50 mL) and DMSO (10 mL). The resulting solids were collected via vacuum filtration and dried to afford (*S*)-1-(methylsulfonyl)-*N*-(4-(methylthio)-1-oxo-1-((4-(3-(pyridin-4-yl)phenyl)thiazol-2-yl)amino)butan-2-yl)-1*H*-pyrrole-3-carboxamide (2.05 g, 77.0% yield) as a white solid. LCMS (ESI) m/z [M + H]^+^ = 555.9; ^1^H NMR (400 MHz, DMSO-d_6_) d 12.49 (s, 1H), 8.68–8.66 (m, 2H), 8.46 (d, J = 7.2 Hz, 1H), 8.31–8.30 (m, 1H), 8.02–8.00 (m, 1H), 7.94–7.96 (m, 1H), 7.83 (s, 1H), 7.73–7.74 (m, 3H), 7.61–7.57 (m, 1H), 7.31–7.29 (m, 1H), 6.79–6.77 (m, 1H), 4.74–4.69 (m, 1H), 3.57 (s, 3H), 2.67–2.53 (m, 2H), 2.13–2.01 (m, 5H). LCMS (ESI) m/z: = 555.9 (M + H^+^).

**Preparation of *N*-[(1*S*)-1-(methoxymethyl)-2-[[4-[3-(1-methylpyrazol-3-yl)phenyl]thiazol-2-yl]amino]-2-oxo-ethyl]-1- methylsulfonyl-pyrrole-3-carboxamide (FHT-2344, Compound 3)**

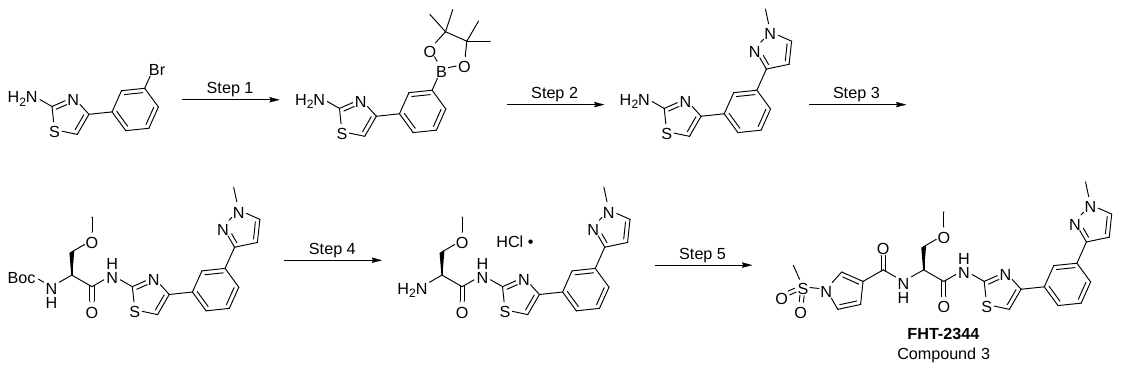

*Step 1. Preparation of 4-(3-(4,4,5,5-tetramethyl-1,3,2-dioxaborolan-2-yl)phenyl)thiazol-2-amine*

To a solution of 4-(3-bromophenyl)thiazol-2-amine (20.0 g, 78.4 mmol), 4,4,5,5-tetramethyl-2-(4,4,5,5-tetramethyl-1,3,2-dioxaborolan-2-yl)-1,3,2-dioxaborolane (21.9 g, 86.2 mmol) and Pd(dppf)Cl_2_ (2 g, 2.73 mmol) in 1,4-dioxane (300 mL) was added KOAc (23.1 g, 235 mmol). After stirring at 80 °C for 2 h, the reaction mixture was cooled to ambient temperatures and concentrated under reduced pressure. The mixture was diluted with water and extracted with three times with ethyl acetate. The combined organic layers were washed with brine, dried with anhydrous Na_2_SO_4_, filtered and concentrated under reduced pressure to afford crude 4-(3-(4,4,5,5-tetramethyl-1,3,2-dioxaborolan-2-yl)phenyl)thiazol-2-amine (23.69 g) as brown solid, which was used to the next step without further purification. LCMS (ESI) m/z: = 303.2 (M + H^+^).

*Step 2. Preparation of 4-(3-(1-methyl-1H-pyrazol-3-yl)phenyl)thiazol-2-amine*

To a solution of 4-(3-(4,4,5,5-tetramethyl-1,3,2-dioxaborolan-2-yl)phenyl)thiazol-2-amine (23.0 g, 76.1 mmol), 3-bromo-1-methyl-pyrazole (12.3 g, 76.1 mmol) and Pd(dppf)Cl_2_ (3.00 g, 4.60 mmol) in a mixture of 1,4-dioxane (200 mL) and water (50 mL) was added K_3_PO_4_ (48.5 g, 228 mmol). After the suspension was stirred at 80 °C for 4 h, the reaction mixture was subsequently cooled to ambient temperatures and concentrated under reduced pressure. The residue was purified by silica gel flash chromatography to afford 4-(3-(1-methyl-1H-pyrazol-3-yl)phenyl)thiazol-2-amine (15.0 g, 76.9% yield) as a gray solid. ^1^H NMR (400 MHz, DMSO-d_6_) d 8.25–8.24 (m, 1H), 7.73–7.63 (m, 3H), 7.38–7.34 (m, 1H), 7.08–6.06 (m, 3H), 6.69–6.68 (m, 1H), 3.89 (s, 3H). LCMS (ESI) m/z: = 257.1 (M + H^+^).

*Step 3. Preparation of tert-butyl N-[(1S)-1-(methoxymethyl)-2-[[4-[3-(1-methylpyrazol-3-yl)phenyl]thiazol-2-yl]amino]-2-oxoethyl] carbamate*

To a solution of 4-[3-(1-methylpyrazol-3-yl)phenyl]thiazol-2-amine (400 mg, 1.56 mmol) and (2*S*)-2-(tertbutoxycarbonylamino)-3-methoxy-propanoic acid (411 mg, 1.87 mmol) in dichloromethane (4.0 mL) was added 2-ethoxy-1-ethoxycarbonyl-1,2-dihydroquinoline (579 mg, 2.34 mmol). The mixture was stirred at 25 °C for 2 h, and then diluted with water. The aqueous layer was adjusted to pH ~6 with 1 N aqueous HCl and the mixture was extracted with twice with ethyl acetate. The combined organic layers were washed twice with saturated aqueous NaHCO_3_ solution, dried with anhydrous Na_2_SO_4_, filtered and concentrated under reduced pressure. The residue was purified by silica gel flash chromatography to afford tert-butyl *N*-[(1*S*)-1-(methoxymethyl)-2-[[4-[3-(1-methylpyrazol-3-yl)phenyl]thiazol-2-yl]amino]-2-oxoethyl] carbamate (0.350 g, 48.4% yield) as a white solid. ^1^H NMR (400 MHz, DMSO-d_6_) d 12.45 (s, 1H), 8.39 (s, 1H), 7.84–7.78 (m, 1H), 7.75 (d, J = 2.0 Hz, 1H), 7.73–7.70 (m, 2H), 7.45–7.43 (m, 1H), 7.15 (d, J = 7.6 Hz, 1H), 6.73 (d, J = 2.0 Hz, 1H), 4.54–4.48 (m, 1H), 3.90 (s, 3H), 3.57 (d, J = 6.0 Hz, 2H), 3.27 (s, 3H), 1.39 (s, 9H). LCMS (ESI) m/z: = 458.2 (M + H^+^).

*Step 4. Preparation of (S)-2-amino-3-methoxy-N-(4-(3-(1-methyl-1H-pyrazol-3-yl)phenyl)thiazol-2-yl)propanamide hydrochloride*

To a solution of tert-butyl *N*-[(1*S*)-1-(methoxymethyl)-2-[[4-[3-(1-methylpyrazol-3-yl)phenyl]thiazol-2-yl]amino]-2-oxoethyl] carbamate (250 mg, 0.546 mmol) in 1,4-dioxane was added a solution of 4 M HCl in 1,4-dioxane (3 mL, 12 mmol). The resulting reaction mixture was stirred at ambient temperatures for 0.5 h and subsequently concentrated under reduced pressure to afford (2*S*)-2-amino-3-methoxy-*N*-[4-[3-(1-methylpyrazol-3-yl)phenyl]thiazol-2-yl]propanamide (215 mg) as a white solid which were used in the next step without further purification. LCMS (ESI) m/z: = 358.1 (M + H^+^).

*Step 5. Preparation of N-[(1S)-1-(methoxymethyl)-2-[[4-[3-(1-methylpyrazol-3-yl)phenyl]thiazol-2-yl]amino]-2-oxo-ethyl]-1-methylsulfonyl-pyrrole-3-carboxamide (****FHT-2344****)*

A solution of (S)-2-amino-3-methoxy-N-(4-(3-(1-methyl-1H-pyrazol-3-yl)phenyl)thiazol-2-yl)propanamide hydrochloride (107 mg, 0.272 mmol), 1-(methylsulfonyl)-1H-pyrrole-3-carboxylic acid (51 mg, 0.272 mmol), 1-hydroxybenzotriazole (110 mg, 0.815 mmol), *N*-(3-dimethylaminopropyl)-*N*′-ethylcarbodiimide hydrochloride (156 mg, 0.815 mmol) and *N*,*N*-diisopropylethylamine (0.142 mL, 0.815 mmol) in dichloromethane (1.0 mL) was stirred at 25 °C for 2 h. The mixture was subsequently diluted with water and extracted twice with ethyl acetate. The combined organic layers were washed twice with brine, dried with anhydrous Na_2_SO_4_, filtered and concentrated under reduced pressure. The residue was purified by reversed-phase prep-HPLC to give *N*-[(1*S*)-1-(methoxymethyl)-2-[[4-[3-(1-methylpyrazol-3-yl)phenyl]thiazol-2-yl]amino]-2-oxo-ethyl]-1-methylsulfonyl-pyrrole-3-carboxamide (57 mg, 38.0% yield) as a white solid. ^1^H NMR (400 MHz, CDCl_3_) d 8.25 (s, 1H), 7.81–7.79 (m, 1H), 7.72–7.70 (m, 2H), 7.42–7.40 (m, 2H), 7.21–7.12 (m, 2H), 7.15–7.10 (m, 1H), 6.75–6.74 (m, 1H), 6.59 (d, J = 2.0 Hz, 1H), 5.10–5.05 (m, 1H), 4.06–4.03 (m, 1H), 3.98 (s, 3H), 3.72–3.68 (m, 1H), 3.45 (s, 3H), 3.20 (s, 3H). LCMS (ESI) m/z: = 529.4 (M + H^+^).

**Synthesis of BRM014 (or 1-(3-(Difluoromethyl)isothiazol-5-yl)-3-(2-fluoro-5-(hydroxymethyl)pyridin-4-yl)urea)**

BRM014 was synthesized according to previously described methods.^1^

**Selectivity Screen**

Compound selectivity was profiled against ATPases and kinases using the KiNativ platform (ActivX). The KiNativ platform is based on a chemoproteomics technology that detects compound binding to ATPases and kinases in biological samples. In this assay, biotinylated acryl phosphates of ATP are added to the cell lysates which irreversibly react and cross-link to conserved lysine residues in the ATP-binding pocket of kinases and ATPases. Mass spectrometry is then used to quantify the binding of ATP to detected kinases and ATPases. In brief, A375 cells were treated with FHT-2344 at indicated concentrations, lysed and followed by addition of the ATP probe in excess. In this way the assay measures the competition between FHT-2344 and the ATP probe in binding to the listed ATPases in cells. The data in Supplementary Fig. 1a is % inhibition of ATP binding to the indicated ATPases. In this assay, a % inhibition >35% is the cut-off for significant inhibition.

**Cell culture and compounds**

Cell lines were purchased from ATCC (MV411, 22RV1, RD, A375, DU4475, HCT116, MP38, MP41, MP46, NCIH1299, LNCAP), Millipore-Sigma (92-1, MEL202). Cell lines were grown according to the ATCC or Millipore-Sigma recommendations, and were tested routinely to ensure that they were negative for mycoplasma. For commercially available compounds: IDE196 (also called LXS196) was purchased from MedChem Express. Paclitaxel, Vorinostat, and Selumetinib were purchased from Selleck Chemicals. Dexamethasone was purchased from Millipore-Sigma.

**Cell proliferation and apoptosis assays**

For 3- and 7-day proliferation assays, LNCAP, NCIH1299, 92-1, MP38, MP41, and MP46 cells were incubated at 37°C with a dose titration of compounds in 96 well plates for 3 days. On day 3, cells were split, and a portion of cells were incubated with Cell-Titer Glo 2.0 (Promega) to measure relative growth on day 3 using an Envision plate-reader (Perkin-Elmer). Remaining cells were re-plated back to the starting density in fresh media and fresh compound, and incubated at 37°C for an additional 4 days, after which relative growth on day 7 was measured as on day 3. Growth relative to a DMSO-control was measured, and curves were fit using a 4-point non-linear regression model, and absolute IC_50_ values were measured in GraphPad Prism. Using these same methods, the following cell lines were tested at ChemPartner: HT1197 (bladder), SKES1, CADDES1, RDES (bone), DU4475, MDAMB415, BT474, HCC1419, CAMA1, BT549, MCF7 (breast), KYSE150 (esophagus), WENRIRB1 (eye), JEKO1, TALL1, RAMOS2G64C10, MM1S, REH, OPM2, LP1, KASUMI1, WSUDLCL2, SKM1, SUPB15, RS411, SUDHL6, PFEIFFER, SUDHL4, REC1, OCIAML5, JURKAT, K562, DAUDI, DB, MEG01 (hematologic), G401, G402 (kidney), PLCPRF5, HLE, HLF (kidney), A549, SW1573, NCIH838, NCIH1299, SBC5, NCIH2444 (lung), COLO829, SKMEL3, SKMEL28, SH4, WM115, A375 (melanoma), SJRH30, A204, RD, HS729 (muscle), TYKNU, OV7 (ovary), 22RV1, PC3, DU145 (prostate).

For apoptosis assays, live cells were incubated with CellEvent Caspase 3/7 Detection Reagent (Life Technologies) or the Incucyte Annexin V Green Dye (Sartorius), and fluorescence was measured in real-time using an Incucyte Zoom (Sartorius).

For SOX10 rescue studies, 92-1 cells were transduced with lentiviral particles expressing empty vector (pLenti-C-Myc-DDK-P2A-Puro) or pLenti-SOX10-Myc-DDK-P2A-Puro (Origene). Seven days post-transduction, cells were incubated with a dose-titration of FHT-1015 or IDE196 and relative growth was measured after 3 days by Cell-Titer Glo (Promega).

**Flow cytometry**

92-1 cells were treated with 100 nM FHT-1015 or 1 µM IDE196. After 24, 48, or 72 hours, cells were harvested and fixed overnight in 70% ethanol. The next day, cells were washed with PBS (ThermoFisher), and resuspended in buffer containing 50 µg/mL propidium iodide (Sigma), 0.1% TritonX-100 (Sigma), and 100 µg/mL RNase A (QIAGEN) in 1x PBS. Fixed and stained cells were incubated in the dark for 20 minutes at 37°C, then cell cycle profiles were measured using a CytoFLEX flow cytometer (Beckman Coulter). Analysis was performed using CytExpert software (Beckman Coulter).

**Quantitative PCR**

Following cell treatments, RNA was isolated using the RNeasy Mini Kit (QIAGEN). Purified RNA (up to 1 µg) was converted to cDNA using the SuperScript IV VILO kit (ThermoFisher). cDNA was diluted 6x in nuclease-free water (Ambion) and 3 µl was inputted per 20 µl qPCR reaction in a 384-well plate with Taqman Fast Advanced master mix (ThermoFisher). Taqman assays used were: Hs01060665_g1 (ACTB), Hs00366918_m1 (SOX10), Hs01117294_m1 (MITF), Hs00181829_m1 (FGF9). qPCR reactions were run using a CFX384 (Bio-Rad), and relative expression was calculated using the 2^-ΔΔCt^ method.

**Western blot**

Following treatments, 92-1 or MP41 cells were harvested, washed in 1x PBS, and lysed directly in 2x NuPAGE LDS sample buffer (ThermoFisher). Samples were resolved on 4-12% Bis-Tris gels (ThermoFisher). Gels were transferred to nitrocellulose membranes (Bio-Rad) using a Trans-Blot semi-dry transfer system (Bio-Rad). Membranes were incubated with antibodies targeting β-Actin, SOX10, and MITF (Cell Signaling). Following incubation with IRDye800CW and IRDye680RD secondary antibodies (LI-COR), relative protein levels were measured on an Odyssey CLx Imaging System (LI-COR).

For SOX10 rescue experiments, 92-1 cells were harvested 7 days post-transduction, lysed in sample buffer (RIPA buffer, 1X NuPAGE LDS sample buffer, 1X NuPAGE sample reducing agent, 1X Halt protease and phosphatase inhibitor cocktail) and heated at 95°C for 5 minutes. Samples were resolved on 4-20% Criterion TGX precast gels (Bio-Rad), and proteins were transferred onto nitrocellulose membrane using the iBlot 2 dry blotting system (Life Technologies). Membranes were probed with antibodies targeting SOX10 and β-Actin (Cell Signaling) as described above.

**RNA-seq and GSEA**

Total RNA was extracted from frozen cell pellets previously washed with PBS. 1 μg RNA per sample was used as input material. Sequencing libraries were generated using NEBNext® Ultra™ RNA Library Prep Kit for Illumina® (NEB, USA) following manufacturer’s recommendations, index codes were added to each sample. mRNA was purified from total RNA using poly-T oligo-attached magnetic beads. Fragmentation was carried out using divalent cations under elevated temperature in NEBNext First Strand Synthesis Reaction Buffer (5X). First strand cDNA was synthesized using random hexamer primer and M-MuLV Reverse Transcriptase (RNase H-). Second strand cDNA synthesis was subsequently performed using DNA Polymerase I and RNase H. Remaining overhangs were converted into blunt ends via exonuclease/polymerase activities. After adenylation of 3’ ends of DNA fragments, NEBNext Adaptor with hairpin loop structure were ligated to prepare for hybridization. In order to select cDNA fragments of preferentially 150~200 bp in length, the library fragments were purified with AMPure XP system (Beckman Coulter). Then 3 μl USER Enzyme (NEB) was used with size-selected, adaptor-ligated cDNA at 37 °C for 15 min followed by 5 min at 95 °C before PCR. Then PCR was performed with Phusion High-Fidelity DNA polymerase, Universal PCR primers and Index (X) Primer. At last, PCR products were purified (AMPure XP system) and library quality was assessed on the Agilent Bioanalyzer 2100 system. Sequencing reads were aligned to version hg38 of the human genome using gencode v30 gene annotations. STAR aligner version 2.0.2 with the following parameters: twopassMode Basic, outFilterMultimapNmax 20, alignSJoverhangMin 8 , alignSJDBoverhangMin 1, outFilterMismatchNmax 999, outFilterMismatchNoverLmax 0.1, alignIntronMin 20, alignIntronMax 1000000, alignMatesGapMax 1000000, outFilterType BySJout, outFilterScoreMinOverLread 0.33, outFilterMatchNminOverLread 0.33, limitSjdbInsertNsj 1200000, outSAMstrandField intronMotif, outFilterIntronMotifs None, alignSoftClipAtReferenceEnds Yes, quantMode TranscriptomeSAM GeneCounts, outSAMtype BAM Unsorted, outSAMunmapped Within, chimSegmentMin 15, chimJunctionOverhangMin 15, chimOutType Junctions WithinBAM SoftClip, chimMainSegmentMultNmax 1, outSAMattributes NH HI AS nM NM choutSAMattrRGline ID:rg1 SM:sm1. Differential gene expression was calculated using limma-voom R package. Geneset enrichment analysis and Gene ontology term overrepresentation analysis was performed using clusterProfiler and DOSE R packages.

**ATAC-seq**

ATAC-seq was performed essentially as previously described.^2^ Following library generation, left- and right- size selection was performed with SPRIselect beads (Beckman Coulter) according to the manufacturer’s recommendations. Sequencing was performed using Illumina HiSeq3 platform and approximately 60M of 150 bp paired end reads were obtained per sample. Sequence analysis was performed as follows. Reads were trimmed for adapter sequences using cutdapt^3^ and aligned to version hg38 of the human genome using bowtie2 (parameters -k 4, -X2000, --local ,--mm).^4^ Multimapping alignments were removed and only the the alignment with the highest score for each read pair was kept. Duplicate reads were removed using Picard Tookit and alignments mapping to mitochondrial genome were excluded.^5^ Alignments were filtered for size smaller than 120 bp and read ends shifted +4 or -5 nucleotides depending on the direction of the read. Peak calling was performed using MACS2.0 with a shift parameter of -75 and an extension size of 150 without modelling.^6^ Signal tracks were created using the fold enrichment option. Consistent peaks among replicates were obtained using IDR using a soft idr threshold of 0.1. The consensus peak list considered was the one containing the largest number of regions after pairwise IDR analysis of all replicates. Differential genome accessibility calculation was performed using limma package. Motif enrichment was performed using Meme suite tools (MEME-chip)^7^ and CISBP transcription motif data from MEME motif database version 12.19. Ridge regression analysis was performed using glmnet R package with coefficients calculated using the minimum estimated lambda. Genomic context annotation was performed using the annotatePeaks function of HOMER package using hg38 version of the human genome.^8^

**ChIP-seq and Enhancer calling**

Treated 92-1 cells were fixed with 1/10 volume freshly prepared Formaldehyde solution (37% Formaldehyde [Sigma], 5M NaCl [Sigma], 0.5 M EDTA pH 8.0 [Sigma], 1M HEPES pH 7.9 [Teknova]), and neutralized with 1/20 volumes of 2.5 M glycine (Sigma). Cells were harvested and washed twice with ice-cold PBS (ThermoFisher) containing 0.5% Igepal (Sigma). During the final wash, 1 mM PMSF was included. Cells were pelleted, snap-frozen on dry ice, and shipped to Active Motif for ChIP-seq with the following antibodies: SMARCA4 (Abcam cat# ab110641), SOX10 (Abcam cat# ab155279), MITF (Active Motif cat# 39789), TFAP2A (Santa Cruz cat# sc-12726), H3K27ac (Active Motif cat# 39133). Single end sequencing was performed with read length of 75 bp at a depth of 30M reads, and was performed on the Illumina platform. Reads were aligned using BWA mem with the default parameters. Alignments with a quality lower than 10 were discarded using samtools and duplicated alignments were removed with picard MarkDuplicates tool. Peaks were called with Macs2 callpeak function and fold enrichment coverage tracks were created using the bdgcmp function of the same package. For enhancer/superenhancer calling, broad peaks were called for H3K27Ac alignments using Macs2 callpeak function with –broad parameter and p-value threshold of 1e-9. ROSE was used to identify the enhancers/superenhancers in hg38 version of the human genome using the t parameter set to 2500.^9,10^ Gene mapping of called enhancers/superenhancers was performed using the geneMapper function of the same package.

**PRO-seq**

MP46 uveal melanoma cells were treated with DMSO or 100 nM FHT-1015, and PRO-seq was performed by Arpeggio Biosciences. Normalized directional bigwig coverage tracks were used with dREG for the identification of regions of bidirectional transcription. Differential enrichment of identified bidirectional transcription regions was performed using R limma package using as input normalized coverage data from consolidated dREG calls extended 200bp in each direction.^11^ Motif analysis was performed using HOMER findMotifsGenome function using the fragment size option “given”.

**Whole exome sequencing**

MP41 cells were treated with 100 nM FHT-1015 for about 3 months, and resistant clones were isolated. Whole exome analysis was performed on genomic DNA processed using Agilent SureSelectXT2 Human All Exon V6 Kit and sequenced to 6G of 150bp paired end reads. Reads were aligned with BWA MEM using default options, duplicated alignments were removed using picard MarkDuplitcates. GATK IndelRealigner was used for indel realignment. GATK HaplotypeCaller was used in discovery mode for variant calling using the following parameters: -dontUseSoftClippedBases -stand_call_conf 20.0. Variants were filtered for QD<2, FS>60 and MQ<40. Variants remaining after filtering were annotated using snpEFF.^12^

**ADME**

ADME properties for FHT-2344 are described in Supplementary Fig. 6a. Log D was measured using the shake flask method. pKa was determined by UV metric measurements at pH between 2 – 12. Phosphate buffer and FaSSIF kinetic solubilities were measured at pH 7.4 and 6.5 respectively. Plasma protein binding was calculated from measured protein binding in 10% human and CD-1 mouse plasma. CYP3A4 inhibition was assessed using both midazolam and testosterone as CYP substrates. Compound in vitro activity against human ether-à-go-go-related gene (hERG) potassium channels was assessed in Chinese Hamster Ovary (CHO) cell line using a manual patch-clamp technique. Female Balb-C nude mice were utilized for low dose mouse pharmacokinetics studies with 3 animals per arm. *IV* arm was dosed at steady state at 0.98 mpk with compound formulated in 9:1 PEG-400 : ethanol (0.500 mg/mL) as a clear solution. *PO* arm was dosed at steady state 5.9 mpk with compound formulated in 20% HP-β-CD in water (2.00 mg/mL) as a clear solution.

**In vivo studies**

The *in vivo* efficacy study was performed at Covance (formerly MI Bioresearch, Ann Arbor MI). 6-7 week old, female nude mice (Hsd: Athymic Nude-*Foxn1^nu^*, Envigo), were implanted subcutaneously in the high axilla with 5 million 92-1 uveal melanoma cells in 50% Matrigel. Upon reaching a mean tumor volume of 186 mm^3^, animals were dosed daily with vehicle (20% HP-β-CD in water) or with FHT-2344 by oral gavage. Tumor volumes and body weights were monitored 3 times per week for 21 days. For pharmacodynamic/pharmacokinetic (PK/PD) studies, animals were dosed daily for two days, and were sacrificed at the indicated timepoints following the last dose. Tumors were collected in RNAlater, homogenized, and qRT-PCR was performed to measure *SOX10* and *FGF9* expression as described above. Plasma was collected at the time of animal sacrifice, and total plasma concentrations of FHT-2344 were measured by LC/MS/MS. All procedures carried out in this experiment were conducted in compliance with the applicable laws, regulations and guidelines of the National Institutes of Health (NIH) and with the approval of MI Bioresearch’s Animal Care and Use Committee. MI Bioresearch is an AAALAC accredited facility.

**Public Datasets**

Median expression of *SMARCA4, SMARCA2, SOX10,* and *MITF* were obtained from The Cancer Genome Atlas (TCGA). CERES and DEMETER dependency scores were from Project Achilles/depmap.org.^13–15^
